# Discovery of a new class of reversible TEA-domain transcription factor inhibitors with a novel binding mode

**DOI:** 10.1101/2022.05.24.493232

**Authors:** Lu Hu, Yang Sun, Shun Liu, Hannah Erb, Alka Singh, Junhao Mao, Xuelian Luo, Xu Wu

## Abstract

The TEA domain (TEAD) transcription factor forms a transcription co-activation complex with the key downstream effector of the Hippo pathway, YAP/TAZ. TEAD-YAP controls the expression of Hippo-responsive genes involved in cell proliferation, development, and tumorigenesis. Hyperactivation of TEAD-YAP activities is observed in many human cancers, and is associated with cancer cell proliferation, survival and immune evasion. Therefore, targeting the TEAD-YAP complex has emerged as an attractive therapeutic approach. We previously reported that the mammalian TEAD transcription factors (TEAD1-4) possess auto-palmitoylation activities and contain an evolutionarily conserved palmitate-binding pocket (PBP), which allows small molecule modulation. Since then, several reversible and irreversible inhibitors have been reported by binding to PBP. Here, we report a new class of TEAD inhibitors with a novel binding mode. Representative analog TM2 shows potent inhibition of TEAD auto-palmitoylation both *in vitro* and in cells. Surprisingly, the co-crystal structure of the human TEAD2 YAP-binding domain (YBD) in complex with TM2 reveals that TM2 adopts an unexpected binding mode by occupying not only the hydrophobic PBP, but also a new side binding pocket formed by hydrophilic residues. RNA-seq analysis shows that TM2 potently and specifically suppresses TEAD-YAP transcriptional activities. Consistently, TM2 exhibits strong anti-proliferation effects as a single agent or in combination with a MEK inhibitor in YAP-dependent cancer cells. These findings establish TM2 as a promising small molecule inhibitor against TEAD-YAP activities and provide new insights for designing novel TEAD inhibitors with enhanced selectivity and potency.

## Introduction

Yes-associated protein (YAP) and transcriptional coactivator with PDZ-binding motif (TAZ) are the major downstream effectors of the evolutionarily conserved Hippo pathway that controls organ size and tissue homeostasis (Pan, 2007; Yu et al., 2015). Beyond their critical roles in development, accumulating evidence shows that YAP/TAZ hyperactivation is frequently linked to tumorigenesis in a broad range of human cancers (Harvey et al., 2013; Pan, 2010; Zanconato et al., 2016b). Importantly, YAP/TAZ alone cannot interact with DNA, therefore, requires the binding of transcriptional factors TEA/TEF-domain (TEAD1-4 in mammals and Scalloped in *Drosophila)* to regulate the expression of Hippo-responsive genes (Wu et al., 2008; Zhao et al., 2008). The transcriptional targets of the TEAD-YAP/TAZ complex are involved in cell proliferation, cell survival, immune evasion and stemness (Moroishi et al., 2015). However, direct targeting YAP/TAZ with small molecules has been shown to be difficult. Therefore, pharmacological disruption of TEAD-YAP/TAZ has been considered as a promising avenue for cancer therapy (Holden and Cunningham, 2018; Johnson and Halder, 2014; Pobbati and Hong, 2020; Zanconato et al., 2016a).

One such strategy is to directly target TEAD-YAP interface with peptidomimetic inhibitors (Jiao et al., 2014; Zhang et al., 2014; Zhou et al., 2015). For instance, a peptide termed “Super-TDU” was designed to block the TEAD-YAP interaction (Jiao et al., 2014). “Super-TDU” mimics TDU domain ofVGLL4 which competes with YAP/TAZ for TEAD binding, and has been shown to suppress gastric cancer growth. However, peptide-based inhibitors generally suffer from poor cell permeability and pharmacokinetic properties, limiting their therapeutic applications. Since TEAD-YAP binding interface is shallow and spanning a large surface area, it is particularly challenging to optimize small molecules for desired potency.

Previously, we and others discovered that TEAD auto-palmitoylation plays an important role in regulation of TEAD stability and TEAD-YAP binding, and loss of TEAD palmitoylation leads to inhibition of TEAD-YAP transcriptional activities (Chan et al., 2016; Holden et al., 2020). More importantly, structural and biochemical studies illustrated that the lipid chain of palmitate inserts into a highly conserved deep hydrophobic pocket (Chan et al., 2016; Noland et al., 2016), away from TEAD­ YAP interface, which is suitable for small molecule binding and suggests that lipid-binding allosterically regulates TEAD-YAP activities.

Over the past years, targeting TEAD auto-palmitoylation has emerged as an attractive strategy for fighting cancers with aberrant YAP activation. To date, several companies and academic research groups have developed small molecule inhibitors against TEAD-YAP activities. A non-steroidal anti­ inflammatory drug, flufenamic acid (FA), has been shown to bind to the lipid-binding pocket of TEAD (Pobbati et al., 2015). Although FA lacks potency to block TEAD function, it demonstrates that the lipid-binding pocket could indeed accommodate small molecule binding. Ever since then, FA scaffold has been extensively explored by medicinal chemists to design TEAD inhibitors, including irreversible inhibitors TED-347 (Bum-Erdene et al., 2019), DC-TEADin02 (Lu et al., 2019), MYF-01-037 (Kurppa et al., 2020), K975 (Kaneda et al., 2020) as well as reversible inhibitor VT103 (Tracy T. Tang et al., 2021). In comparison, non-FA based TEAD inhibitors are relatively limited, and only a few examples, such as compound 2, have been reported (Holden et al., 2020). Among the reported inhibitors, K975 and VT103 showed strong anti-proliferation effects *in vitro* and anti-tumor effects *in vivo*. However, these inhibitors only have effects in limited cell lines, such as NF2-defecient mesothelioma cells. In addition, most of the reported TEAD inhibitors are irreversible inhibitors targeting the cysteine at the palmitoylation site, which might have undesired non-specific reactivity towards other cysteines or other targets. To gain insights into the chemical diversity of reversible TEAD inhibitors and their utilities in cancer therapeutics, it is important to identify new chemical scaffolds to target TEADs.

We previously developed a non-FA based reversible TEAD inhibitor, MGH-CP1 (Li et al., 2020a), which inhibited transcriptional output of TEAD-YAP *in vitro* and *in vivo*. However, MGH-CP1 only showed sub-micromolar potency against TEAD palmitoylation *in vitro* and was used at low micromolar range in cellular assays. These limitations prompt us to develop new TEAD inhibitors with higher potency. In this study, we discovered a series of novel TEAD inhibitors featuring a common 4-benzoyl­ piperazine-1-carboxamide scaffold. Among them, TM2 exhibits strong inhibition of TEAD2 and TEAD4 auto-palmitoylation *in vitro* with the ICSO values of 156 nM and 38 nM, respectively. In addition, palmitoylation of both exogenous Myc-TEAD1 and endogenous Pan-TEADs is also significantly diminished by TM2 in HEK293A cells, which further confirms its potency and mode-of­ action in cellular context. The co-crystal structure of TEAD2 YBD in complex with TM2 uncovered a novel binding mode of the compound, which extended into a previously unknown hydrophilic side pocket adjacent to the PBP, and caused extensive side chain rearrangements of the interacting residues. Further functional studies showed that TM2 significantly inhibits YAP-dependent liver organoid growth *ex vivo,* and inhibits proliferation of YAP-dependent cancer cells as a single agent or in combination with a MEK inhibitor. Overall, these studies broaden our understanding of the small molecule binding sites on TEADs.

## Results

### Identification ofTM2 as a novel TEAD auto-pabnitoylation inhibitor

To identify new chemotypes that could inhibit TEAD auto-palmitoylation, we screened a library containing about 30,000 non-proprietary medicinal chemistry compounds with three rounds of click­ ELISA assay (Lanyon-Hogg et al., 2015), through the Astellas-MGH research collaboration by using the recombinant TEAD2 and TEAD4 YBD proteins. The inhibition of ZDHHC2 was used as a selectivity filter *(Figure 1-figure supplement 1)*. We found several hits that share a common 4-(3-(2-cyclohexylethoxy)benzoyl)-piperazine-1-carboxamide moiety (data not shown, with micromolar potency in TEAD palmitoylation assays *in vitro)*. The main variation is located at theN-substituent of the urea moiety with frequent incorporation of heteroarenes. Inspired by this structural convergence, we first designed a series of derivatives with variable substituents at the urea moiety, represented by TM2 and TM22 *(Figure 1A)*. TEAD2 auto-palmitoylation *in vitro* assay was used to evaluate their potency. Compared to heteroaryl group, phenyl substituent showed stronger inhibition on TEAD2 auto­ palmitoylation (TM2 vs. TM22, *Figure 1B)*. Inspired by these results, we explored the tolerance level by increasing hydrophilicity of TM2. As illustrated by TM45 and TM98, hydrophilic groups at the left cyclohexyl ring significantly decrease the activities, while the phenyl moiety at the right-side of the urea moiety is well tolerated (TM112, *Figure 1B)*. Overall, TM2 was identified as the most potent compound *(Figure 1B)* and selected for further biological evaluations.

**Figure 1.**
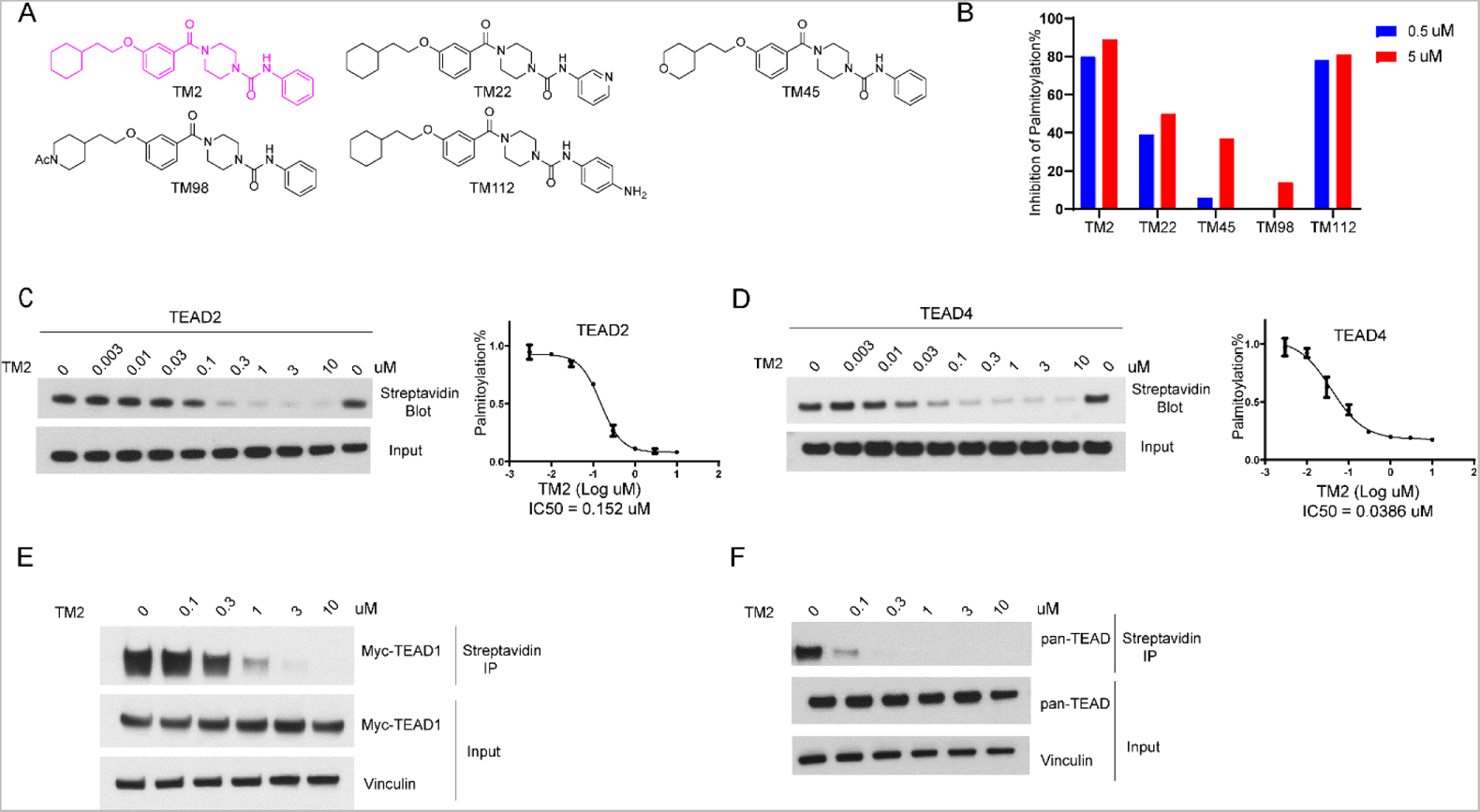
Identification of TM2 and analogues as novel TEAD auto-palmitoylation inhibitors. **(A)** Representative chemical structures of a novel class of TEAD inhibitors with 4-(3-(2-cyclohexylethoxy)benzoyl)-piperazine-1-carboxamide moiety. TM2 structure is highlighted in magenta. **(B)** Inhibition of TEAD2 auto-palmitoylation with treatment of TM2 under 0.05 and 0.5 11M for 30 mins, respectively. IC50 values for TM2 inhibition of TEAD2 **(C)** and TEAD4 **(D)** auto-palmitoylation were characterized by western blot analysis (left) and quantified by Image J (right). The data was determined by independent replicates (n 3), and shown as mean ± SEM. Palmitoylation of Myc­ TEAD1 **(E)** and endogenous pan-TEAD **(F)** were analyzed by immunoprecipitation assay with treatment of TM2 at indicated concentrations for 24 h.

**Figure 1-figure supplement 1.**
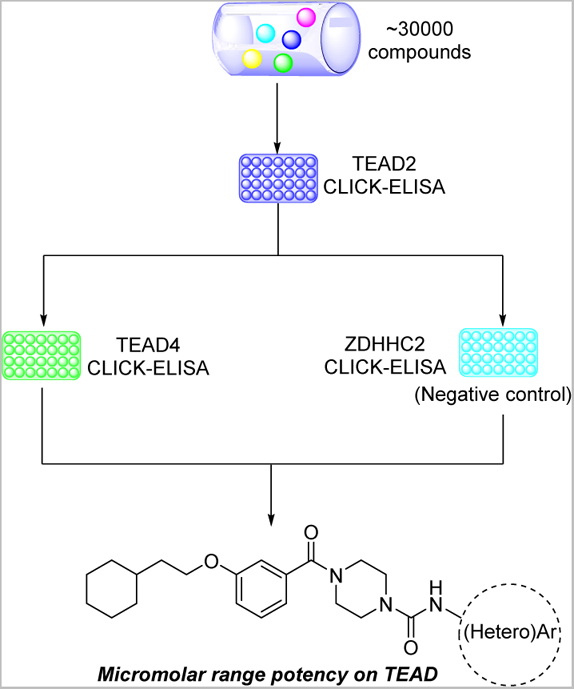
Scheme for High through-put screening of TEAD inhibitors

TEAD family consists of four homologous members, TEAD1-4, which share highly conserved domain architectures (Pobbati and Hong, 2013). We found that TM2 inhibits TEAD2 palmitoylation with an IC_50_ value of 156 nM *(Figure 1C)*. Encouragingly, TM2 displays an even more potent effect on TEAD4 auto-palmitoylation with an IC_50_ of 38 nM *(Figure 1D)*. To study its effects on cellular TEAD palmitoylation, we overexpressed Myc-TEAD1 in HEK293A cells and treated with TM2 at different doses. As **Figure 1E** shows, TM2 dramatically suppresses Myc-TEAD1 palmitoylation in cells in a dose-dependent manner. Furthermore, treatment ofTM2 also significantly inhibits endogenous TEAD1-4 palmitoylation using an antibody recognizing pan-TEADs (***Figure 1F***). Even at as low as 100 nM, Pan-TEAD palmitoylation was diminished. Collectively, these results suggested that TM2 is a potent and pan-inhibitor of palmitoylation of TEAD family proteins.

### TM2 adopts a novel binding mode compared to other known TEAD inhibitors

To gain insights into the precise binding mode of TM2, we determined the co-crystal structure of TEAD2 YBD in complex with TM2 at 2.4 A resolution *(Figure 2, Figure 2-figure supplement 1* **and supplement Table 1**). Overall, TM2 binds to the same PBP in TEAD2 where palmitic acid and other inhibitors target. As shown in **Figure 2B**, the (2-cyclohexylethoxy)phenyl moiety of TM2 is surrounded by several hydrophobic residues, such as F233, L383, L390, F406, 1408, Y426, and F428, enabling strong hydrophobic interactions, which is very similar to the interaction mode of TEAD with the fatty acyl chain of palmitic acid.

**Figure 2.**
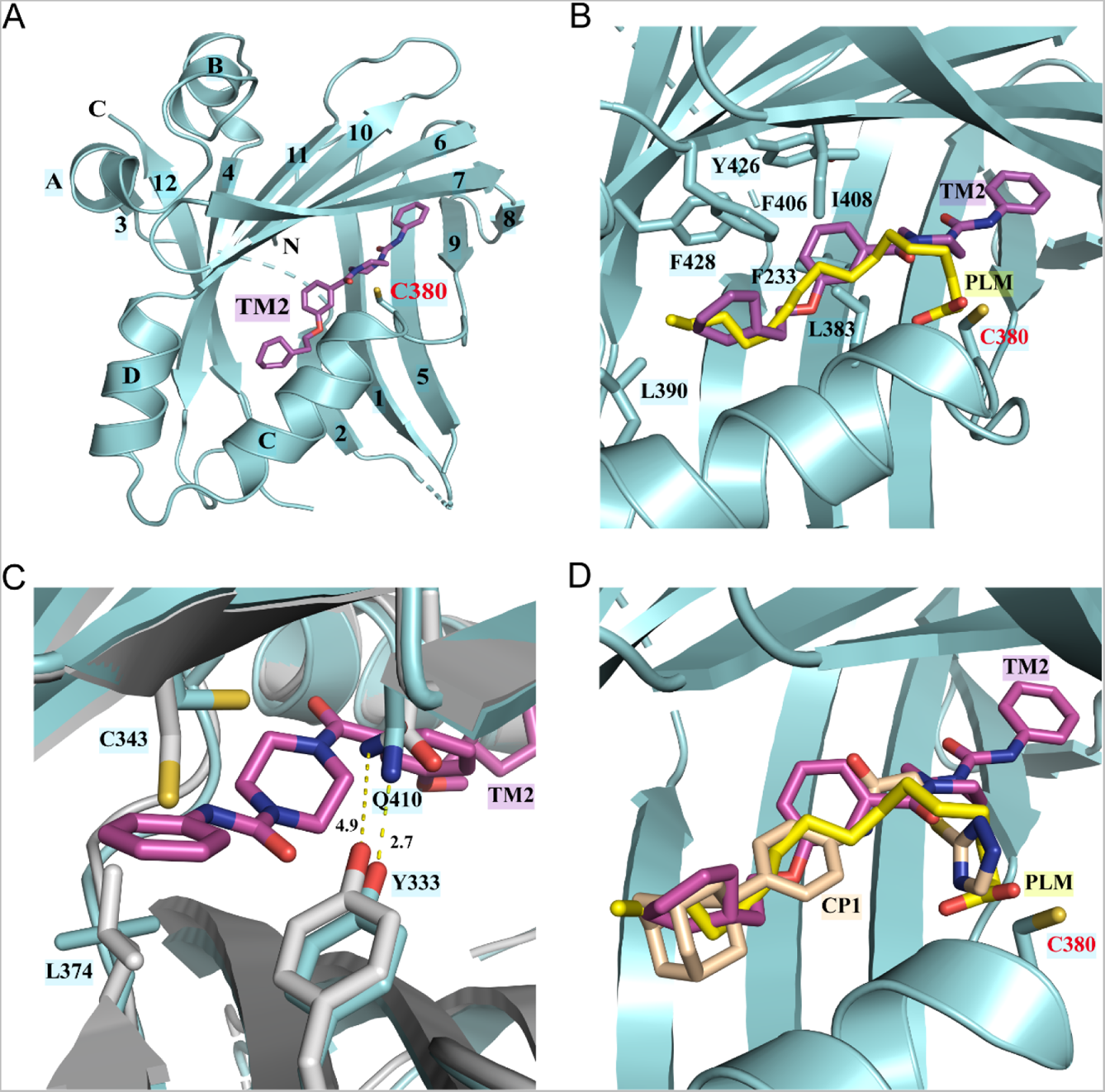
Co-crystal structure of TEAD2 complexed with TM2. (A) Ribbon diagram of the crystal structure of TEAD2-TM2 (PDB 8CUH). TM2 is shown as magenta sticks. (B) Close-up view of the TM2 binding site of TEAD2 (PDB 8CUH) with the superposition of the TEAD2-PLM structure (PDB SHGU). Surrounding residues are shown as cyan sticks. PLM is shown as yellow sticks. (C) Conformational changes in side chains of residues in the new pocket in the presence of TM2 binding. Indicated residues from TEAD2-TM2 and TEAD2-PLM are shown as cyan and gray sticks, respectively. Distances between atoms are shown with yellow dash lines and the unit is angstrom. (D) Structural superposition of TEAD2-TM2 (PDB 8CUH), TEAD2-PLM (PDB SHGU), and TEAD2-CPl(PDB 6CDY). TEAD2 is shown as cyan ribbon. TM2, PLM, and CPl are shown as sticks and colored in magenta, yellow, and wheat, respectively. PLM, Palmitic acid.

**Figure 2-figure supplement 1.**
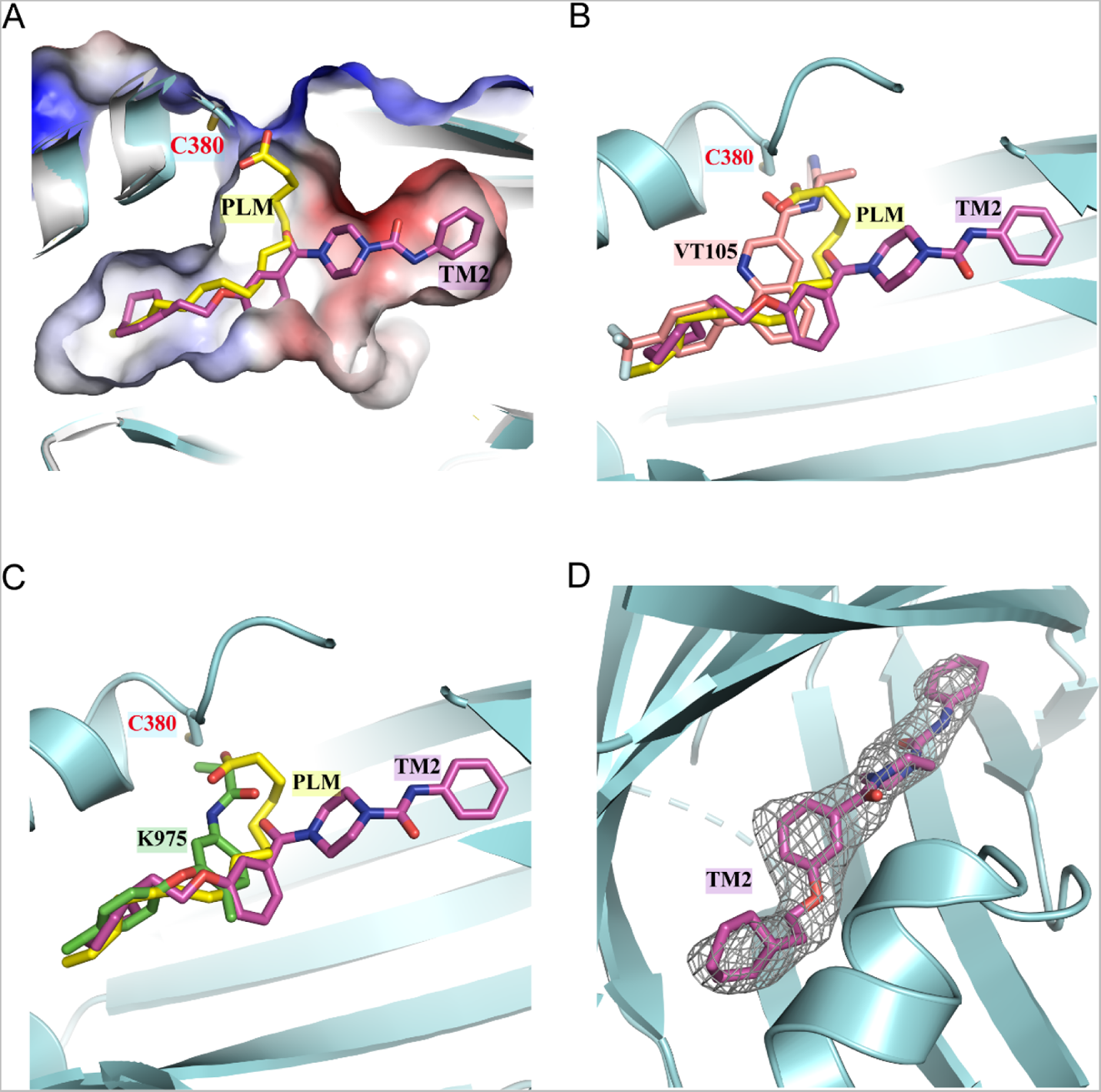
(A) Comparison of orientations of TM2 and PLM in the binding pocket. The TEAD2 protein is shown in cyan ribbon. The pocket is shown by surface. PLM and TM2 are shown as sticks and colored in yellow and magenta, respectively. (B) Structural superposition of TEAD2-TM2 (PDB 8CUH), TEAD2-PLM (PDB SHGU), and TEAD3-VT105 (PDB 7CNL). TM2, PLM, and VT105 are shown as sticks and colored in magenta, yellow, and salmon, respectively. (C) Structural superposition of TEAD2-TM2 (PDB 8CUH), TEAD2-PLM (PDB SHGU), and TEAD1-K975 (PDB 7CMM). TM2, PLM, and K975 are shown as sticks and colored in magenta, yellow, and green, respectively. (D) The *F*_0_ - *F*c omit electron density map for TM2 at the contour level of 2.5 *a* is shown in gray. The TEAD2 protein is shown in cyan ribbon and TM2 is shown as magenta sticks.

**Figure 2-supplement Table 1.**
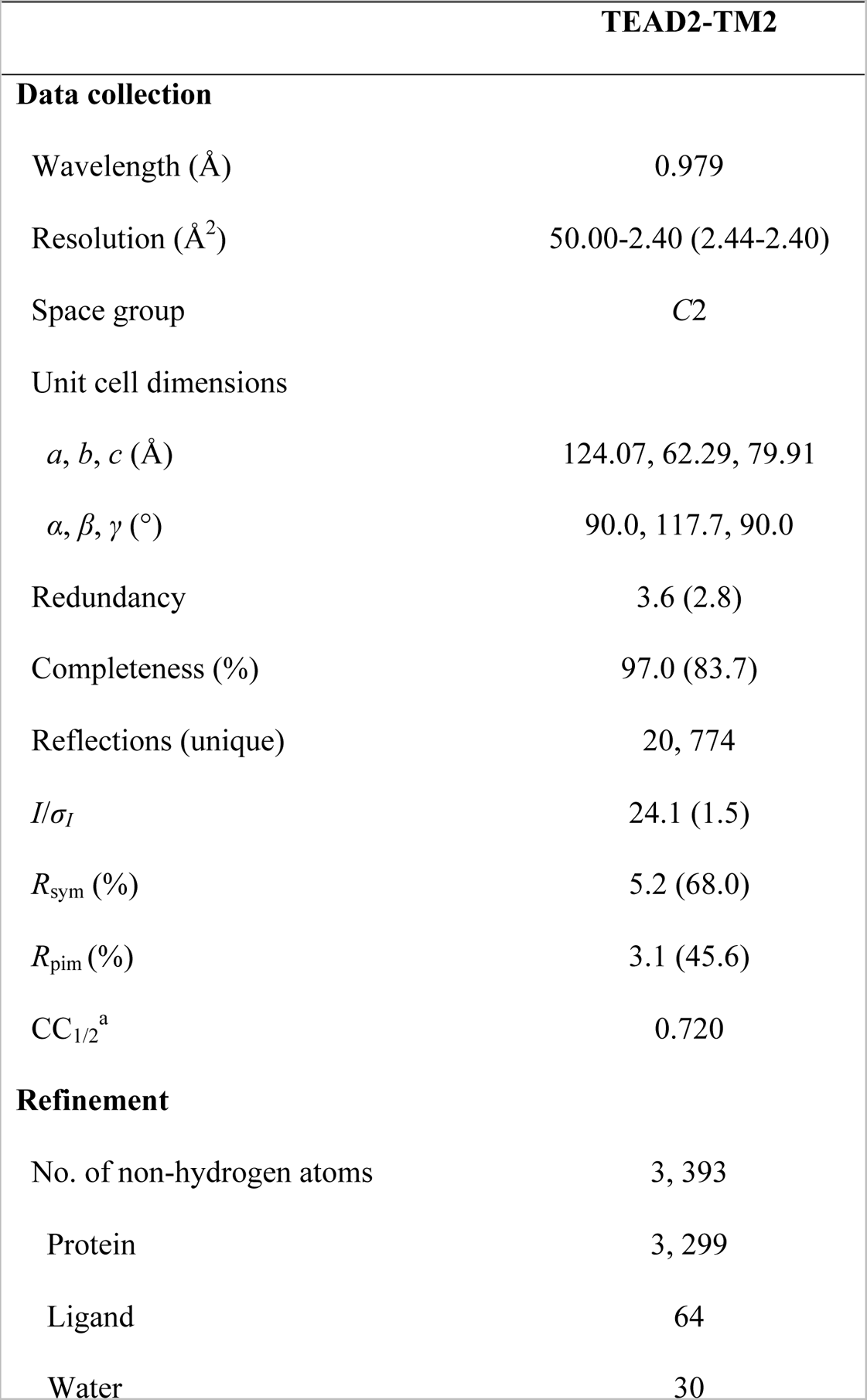
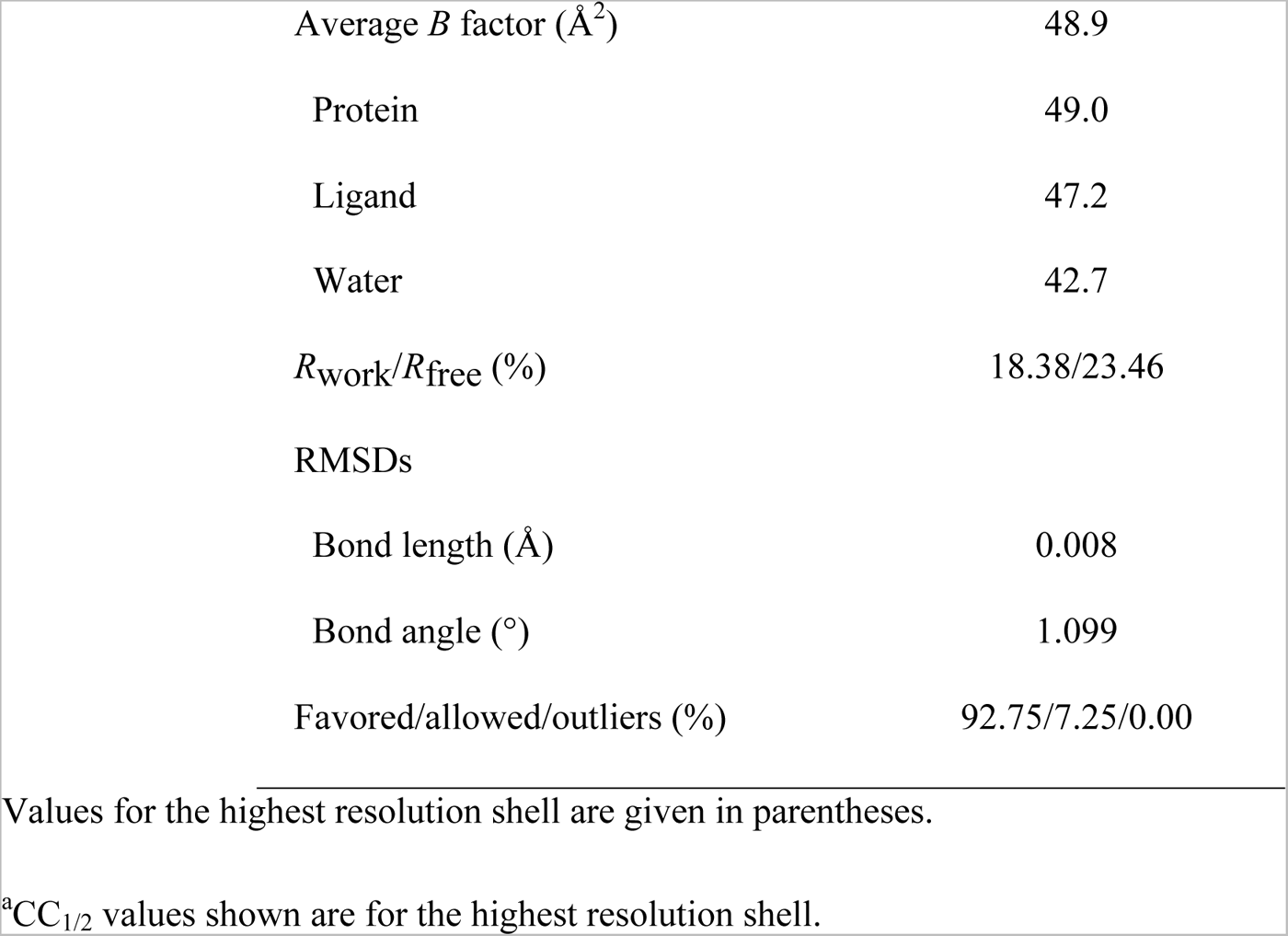
Data collection and structure refinement statistics.

However, by superposing the TEAD2-TM2 (PDB 8CUH) with TEAD2-PLM structures (PDB 5HGU) (Chan et al., 2016), we observed a new feature of TM2 binding *(Figure 2B)*. Unlike palmitic acid with its head group pointing towards residue C380, the urea moiety of TM2 exhibits a completely different orientation and sticks into a new side pocket, which has never been reported before to be involved in TEAD inhibitor binding and is only accessible by rearranging the side chains upon TM2 binding *(Figure 2B and* Figure 2-figure supplement 1A). TM2 binding drives significant conformational changes in the side chains of residues C343 and L374, which makes space for TM2 insertion *(Figure 2C)*. Additionally, TM2 binding causes the side chain movement in residue Q410 and Y333, which reduces the distance between the nitrogen atom ofQ410 and the oxygen atom ofY333 from 4.9 A to 2.7 Å to allow the formation of favorite electrostatic interaction *(Figure 2C)*.

This binding model is highly consistent with our structure-activity relationship (SAR) results in *Figure 1A-B* that demonstrate that the left hydrophobic tail is repulsive to incorporate hydrophilicity, while the urea moiety is tolerated. The surface electrostatics of the TM2 binding pocket (*Figure 2-figure supplement 1A*) also illustrated that the (2-cyclohexylethoxy)phenyl moiety inserts into a nearly neutral environment, while the urea is buried in a pocket bearing electronegative properties. Furthermore, the electronegative carbonyl which links benzene and piperazine is spatially adjacent to electropositive electrostatics.

We then set to figure out whether this unexpected binding model is unique to TM2, compared to other TEAD inhibitors. The co-crystal structures of TEAD YBD in complex with PLM (PDB 5HGU), TM2 (PDB 8CUH), and other known TEAD inhibitors, including MGH-CP1 (PDB 6CDY) (Li et al., 2020a), K975 (PDB 7CMM) (Kaneda et al., 2020) and VT105 (PDB 7CNL) (Tracy T. Tang et al., 2021), were superposed *(Figure 2D* and *Figure 2-figure supplement 1B and 1C)*. Although PLM and these TEAD inhibitors are co-crystallized with different members of TEAD family of proteins, the highly homologous structures of TEAD YBD allowed us to compare their binding modes. Consistent with previously reported results, MGH-CP1, VT105 or K975 adopts almost the same binding mode as PLM and fits very well with the PBP. However, the scenario depicted by TM2 is quite different, which provides new insights into the structural adaptability for development of TEAD inhibitors. Considering relatively higher hydrophilicity in the new side pocket, there will be much more space to balance the lipophilicity of TEAD inhibitors and improve drug-like properties, such as solubility and metabolism (Waring, 2010).

### TM2 inhibits TEAD-YAP association and TEAD-YAP transcriptional activity

TEAD auto-palmitoylation plays important roles in regulation of TEAD-YAP interaction. To confirm whether TM2 functions through blockade of TEAD-YAP binding, we tested TM2 in a malignant pleural mesothelioma (MPM) cell line H226 cells, which is deficient with NF2 and highly dependent on TEAD­ YAP activities (Kaneda et al., 2020; Tracy T Tang et al., 2021). YAP co-immunoprecipitation (IP) experiments indicated that TM2 dramatically blocked the association of YAP with endogenous TEADl as well as pan-TEAD in a dose-dependent manner *(Figure 3A)*. Next, we evaluated the effects of TM2 in the expressions of TEAD-YAP target genes, represented by *CTGF, Cyr61* and *ANKDRJ*. After treatment of TM2, the expression levels of *CTGF* and *ANKDRJ* were significantly suppressed at both 24 and 48 h, while *Cyr61* show strong response at 48h *(Figure 3B and Figure 3-figure supplement 1)*.

**Figure 3.**
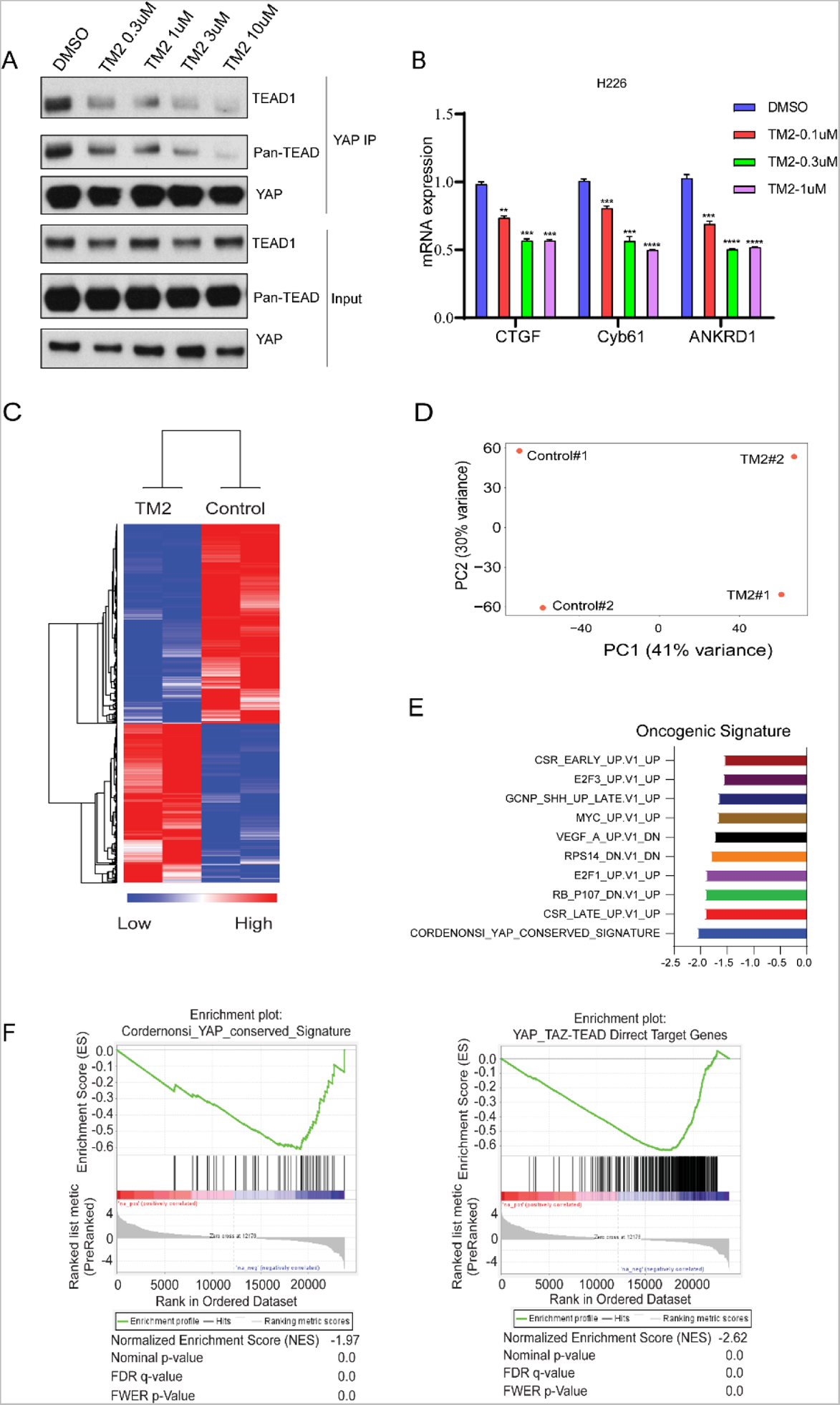
TM2 suppressed transcriptional outputs of Hippo pathway in cancer cells. (A) H226 cells were treated with TM2 at indicated concentrations for 24 h. The interactions of YAP and Pan-TEAD as well as TEADl was observed with YAP Co-IP. **(B)** representative target genes of Hippo pathway in H226 cells were measured with treatment of TM2 at indicated concentrations for 48 h. The data was determined by independent triplicates (n 3) and shown as mean ± SEM. Significance was determined by two-tailed t-test. \*\**P* < 0.01, \*\*\**P* < 0.001, \*\*\*\**P* < 0.0001. **(C)** Heatmap analysis of global genes transcriptional alteration in H226 treated with vehicle control or TM2. **(D)** PCA biplot with genes plotted in two dimensions using their projections onto the first two principal components, and 4 samples (Control 2 samples, TM2 2 samples) plotted using their weights for the components. (E) Gene set enrichment analysis of H226 cells treated with TM2 using oncogenic signature gene sets from Molecular Signatures Database. **(F)** Gene set enrichment plot of Cordernonsi_YAP_conserved_Signature (left panel) and YAP_TAZ-TEAD Direct Target Genes (right panel) with H226 cells treated with TM2.

**Figure 3-figure supplement 1.**
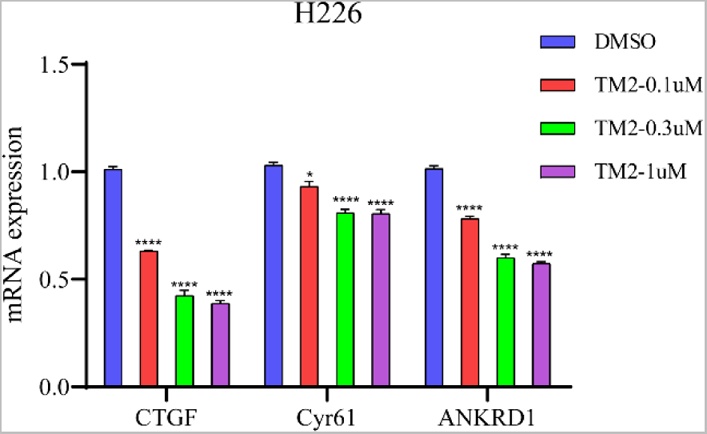
Target gene expression in H226 with TM2 treatment for 24 h. The data was determined by independent triplicates (n 3) and shown as mean ± SEM. Significance was determined by two-tailed t-test. *\*p* < 0.05, *\*\*\*\*p* < 0.0001.

**Figure 3-figure supplement 2.**
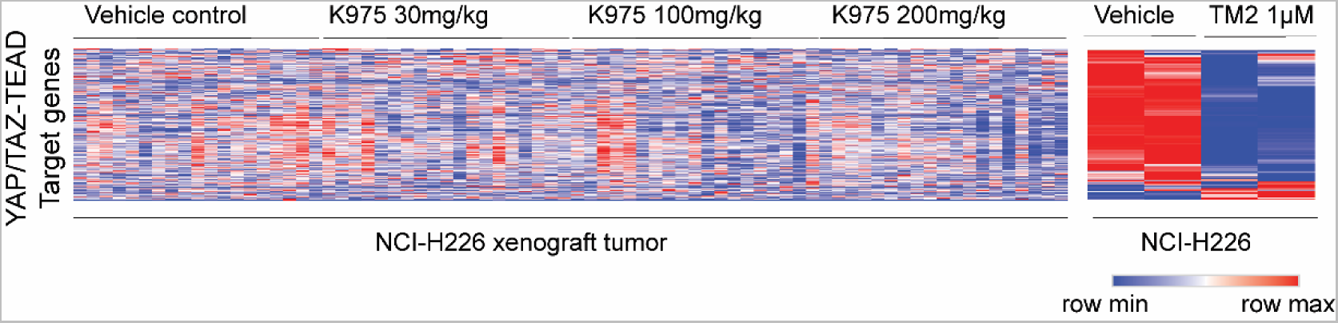
Heatmap analysis ofYAP/TAZ-TEAD direct target genes transcriptional alteration in H226 xenograft tumor treated with K975 (GSE196726) and H226 cell line treated with TM2 (lμM).

In order to systemically evaluate the effect of TM2 on YAP/TAZ-TEAD transcriptional activation, we performed RNA-seq analysis *(Figure 3C)*. YAP/TAZ-dependent H226 cells were treated with or without TM2. We performed principle component analysis (PCA), a mathematical algorithm reducing the dimensionality of the data while retaining most of the variation in the data sets. The samples were plotted and indicated that TM2 treatment substantially altered the gene sets at PC! in H226 cells *(Figure 3D)*. Gene set enrichment analysis (GSEA) was performed to analyze the transcriptional signature gene sets from Molecular Signature Database. It showed that YAP signature was the top enriched signature according to the Normalized Enrichment Score (NES) *(Figure 3E)*. To further validate the effects of TM2 on YAP/TAZ signaling, the Cordernonsi_YAP_conserved_Signature and YAP_TAZ-TEAD Direct Target Genes were determined (Zanconato et al., 2015). Consistently, YAP/TAZ signature was significantly enriched in downregulation phenotype in both of gene sets *(Figure 3F)*. We then compared the specificity of TM2 with that of irreversible TEAD inhibitor K975 which showed strong antitumor effects in H226 xenograft tumor. Through global analysis of YAP/TAZ­ TEAD direct target genes in H226 xenograft tumor treated with three doses of K975 (p.o.) and H226 cells treated with lf!M TM2 (Kaneda et al., 2020; Zanconato et al., 2015), we found that TM2 was more efficient to block YAP/TAZ-TEAD target genes relative to K975 in H226 xenograft tumors *(Figure 3-figure supplement 2),* highlighting the high specificity of our reversible inhibitors. Taken together, we identified TM2 as a potent disruptor that can specifically attenuate outputs of Hippo pathway.

### TM2 inhibits YAP-dependent organoids growth and cancer cell proliferation

YAP activity has been shown to be critical for the growth of liver organoid (Planas-Paz et al., 2019). Therefore, we used mouse hepatic progenitor *ex vivo* organoids to further investigate the effects of TM2 in a physiologically relevant model. As shown in **Figure 4A**, TM2 impaired the sustainability of organoids growth in a dose dependent manner, with more than 85% of disruption at 40 nM. Consistently, Ki67 positive cells for organoids maintenance in 3D culture were significantly diminished upon TM2 treatment *(Figure 4B* and *Figure 4-figure supplement 1)*.

**Figure 4.**
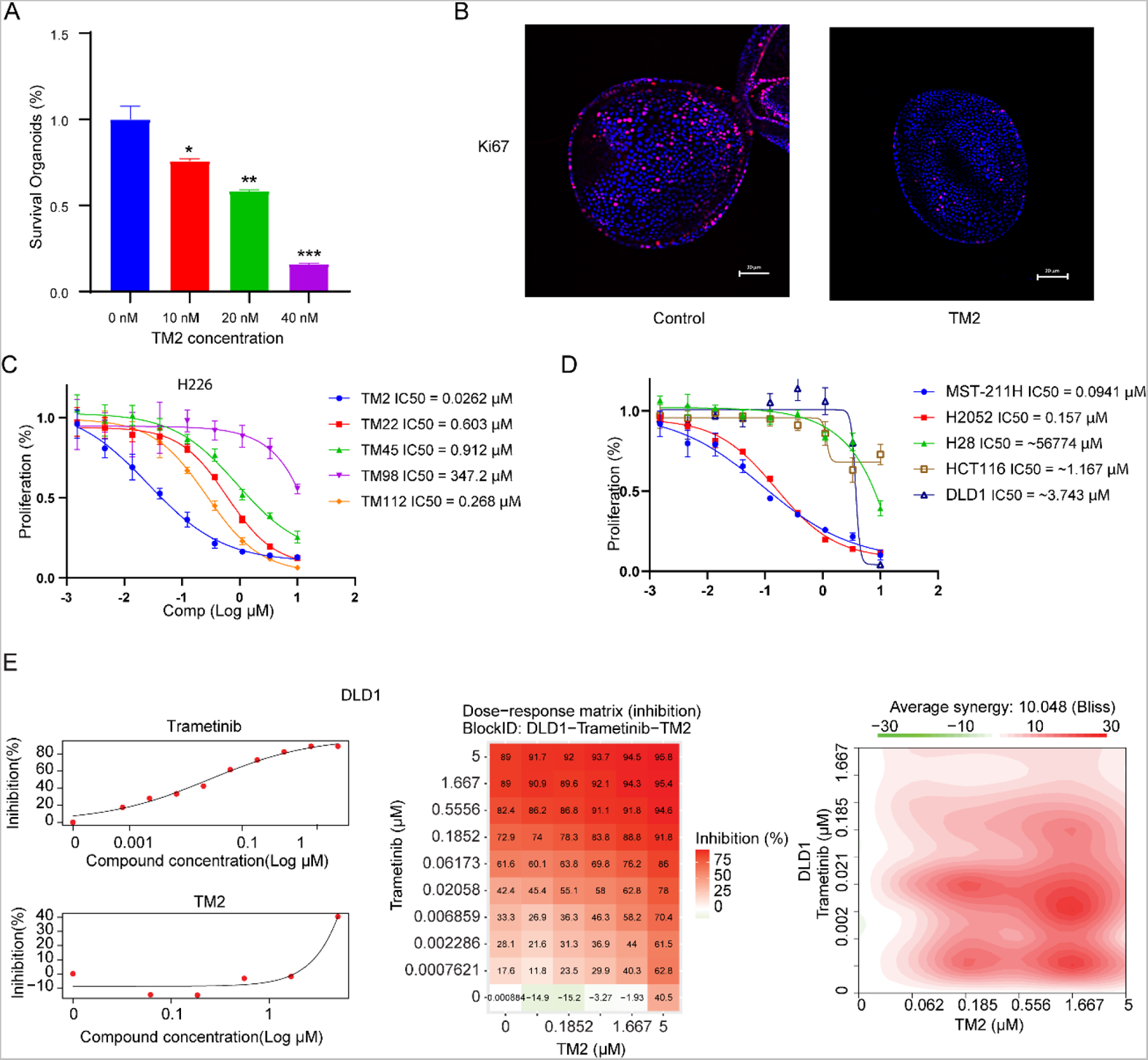
TM2 showed inhibition on YAP dependent proliferation. **(A)** Percentages of survival organoids with treatment of control or TM2 at indicated concentrations. The data was determined by independent triplicates (n=3) and shown as mean± SEM. Significance was determined by two-tailed t-test. \**P* < 0.05, ** *P* < 0.01, \*\*\**P<* 0.001 (B) Immunofluorescent staining of Ki67 in organoids treated with control or TM2 (40 nM). Pink, Ki-67; blue, nuclear DNA (DAPI). Bar, 20 11m. (C) Cell inhibition in H226 cells with treatment of compounds at indicated concentrations for 6 days. The data was determined by independent triplicates (n 3) and shown as mean± SEM. (D) Cell inhibition in MSTO-211H, H2052, H28, HCT116 and DLDl cells with treatment of TM2 at indicated concentrations for 5, 7, 6, 5, or 5 days, respectively. The data was determined by independent triplicates (n=3) and shown as mean± SEM. (E) Drug combination experiments using TM2 and MEK inhibitor Trametinib in DLDl: Heatmaps show color-coding as percentage of cell viability normalized to untreated controls. Heatmaps of Bliss score for TM2 and Trametinib combination were shown.

Pleural mesothelioma (MPM) is a type of aggressive tumor, associated with exposure to asbestos fibers (Rossini et al., 2018). Despite several standard therapies, such as surgery, radiotherapy, chemotherapy and immunotherapies, MPM patients still suffer poor prognosis with a median survival of only 8-14 months (Nicolini et al., 2020). *NF2* and *LATS2,* the upstream components of Hippo pathway, are frequently observed to be inactivated in malignant mesothelioma (MM), leading YAP activation in more than 70% of analyzed primary MM tissues (Murakami et al., 2011; Sekido, 2018). Therefore, MM would be a good model to study the therapeutic effects of TM2 on Hippo signaling defective cancers. Encouraged by the strong inhibition of TEAD-YAP transcriptional activities in H226 cells, we first evaluated anti-proliferative activities of TM2 in this cell line. As shown in **Figure 4C**, H226 cells exhibited striking vulnerability to TM2 treatment with an IC_50_ value of 26 nM, consistent with its potency in blocking TEAD palmitoylation in vitro and in cells. Other derivatives, including TM22, TM45, TM98, TM112 are less potent as TM2, which correlated well with their *in vitro* activities *(Figure 4C)*. In addition, we also studied the effects of TM2 in two other MPM cell lines, MST0-211H and NCI-H2052, which harbors *Lats1/2* deletion/mutations, and NF2-deficiency, respectively (Kaneda et al., 2020; Lin et al., 2017; Miyanaga et al., 2015). Consistently, TM2 also significantly inhibits cell proliferation of MST0-211H and NCI-H2052 cells *(**Figure 4D**)* with IC_50_ values of 94 nM and 157 nM, respectively. In comparison, TM2 shows no significant inhibition in the Hippo WT mesothelioma cells, NCI-H28 with IC_50_ >5 11M (Tanaka et al., 2013) *(**Figure 4D**),* suggesting TM2 is specific to YAP-activated cancer cells.

Currently, TEAD inhibitors mainly show promising therapeutic potentials in mesothelioma, with limited activities in other YAP-dependent cancer cells. Given that deregulated Hippo signaling is implicated in many human cancers (Harvey et al., 2013), it is important to test the efficacy of TEAD inhibitors in cancers beyond mesothelioma, which will deepen our understanding of therapeutic spectrum of blocking TEAD-YAP activities. Therefore, we evaluated TM2 in colorectal cancer (CRC), as Hippo pathway has been shown to regulate the progression of CRC (Della Chiara et al., 2021; Jin et al., 2021; Pan et al., 2018). However, TM2 did not exhibit strong inhibition on cell proliferation of two CRC cell line *(Figure 4D),* HCT116 and DLD1. These results suggested suppression of Hippo transcriptional activities in CRC alone might not be sufficient to inhibit cell growth, as observed in mesothelioma. Indeed, YAP are found to be capable of rescuing cell viability in HCT116 with loss function of KRAS, implying KRAS signaling might also account for lack of potency of TM2 in CRC. Hence, we performed a drug combination matrix analysis across 5 doses of TM2 and 9 doses of MEK inhibitor trametinib in HCT116 and DLD1, respectively. Encouragingly, we observed strong inhibitory effects and substantial synergy in both of two cell lines *(Figure 4E and Figure 4-figure supplement 2),* suggesting that combining TEAD inhibitors with other therapies might be a good strategy to broaden their therapeutic applications in near future. Together, our data highlights that TM2 might have appealing potentials to antagonize carcinogenesis driven by aberrant YAP activities.

**Figure 4-figure supplement 1.**
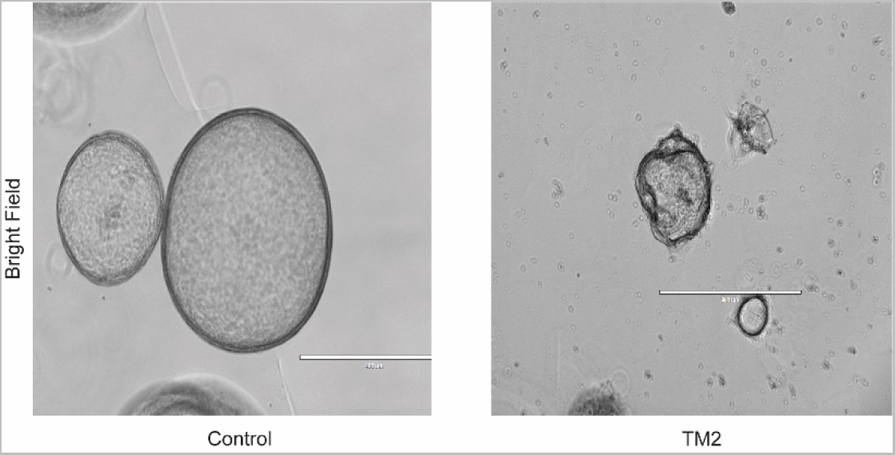
Bright field images of organoids treated with control or TM2 (40 nM). Bar, 400 11m.

**Figure 4-figure supplement 2.**
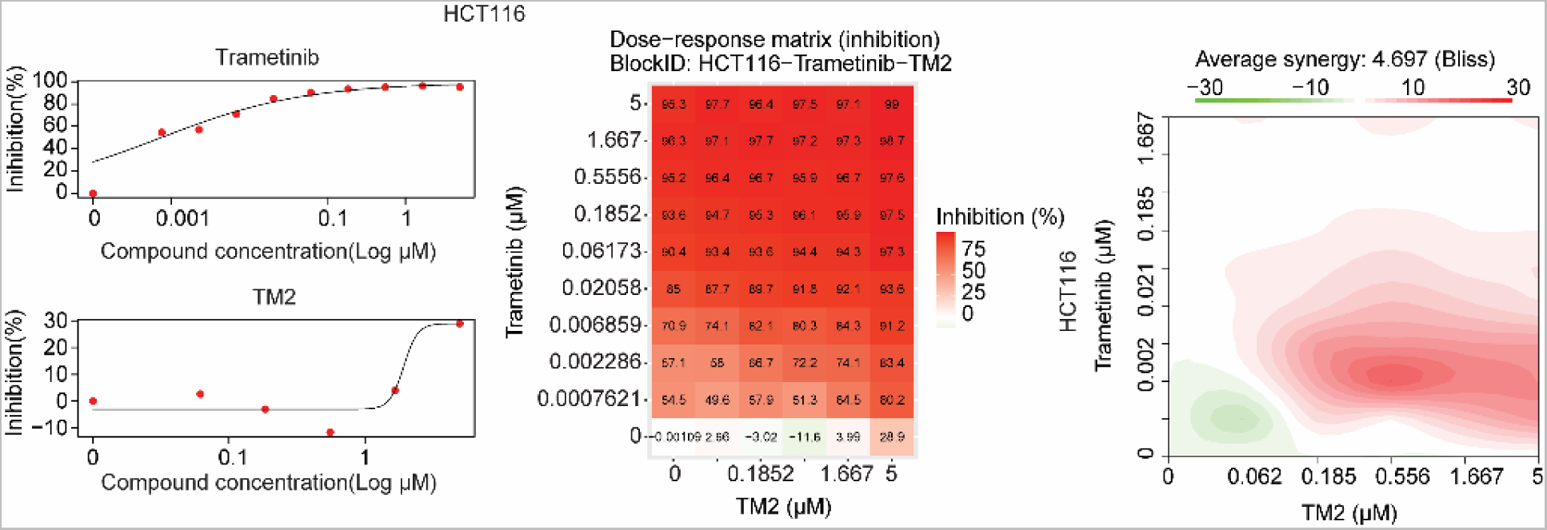
Drug combination experiments usmg TM2 and MEK inhibitor Trametinib in HCT116: Heatmaps show color-coding as percentage of cell viability normalized to untreated controls. Heatmaps of Bliss score for TM2 and Trametinib combination were shown.

## Discussions

In this study, we discovered a new class of reversible pan-TEAD inhibitors. The most potent compound, TM2, significantly diminished TEAD2/4 auto-palmitoylation in nanomolar ranges. Co-crystal structure analysis of TM2 in complex with TEAD2 YBD discovered a novel binding mode. It showed that the much more hydrophobic part of TM2, featured a cyclohexyl ring, exquisitely fits the palmitoylation pocket, which is the most well-known structure feature for targeting TEAD (Dey et al., 2020). Surprisingly, the structure demonstrates that the urea moiety of TM2, does not overlap with PLM, but sticks into a new and unique side pocket. This binding site is not occupied by all other known TEAD inhibitors, such as MGH-CPI, VTIOS and K975. This side pocket is not fully available in the palmitate­ bound TEAD2 structure, but is formed with significant side chain rearrangement upon TM2 binding. Moreover, this side pocket is endowed with higher hydrophilicity than the lipid-binding pocket, providing potentials for enhancing drug-like properties. The novel binding model expands structural diversity of the TEAD binding pocket and will boost the discovery of more novel chemotypes, contributing to the development of therapeutics targeting TEAD-YAP.

Blocking TEAD auto-palmitoylation by TM2 disrupted TEAD-YAP association. Consistently, we observed significant suppression of downstream Hippo transcription program with treatment of TM2. RNA-seq analysis further confirmed that TM2 specifically inhibits YAP transcriptional signatures. YAP/TAZ is constitutively active in many human malignancies and shown to be essential for many cancer hallmarks (Zanconato et al., 2016b). Therefore, targeting YAP/TAZ activities has been considered as an attractive strategy for cancer therapy. In human MPM, a type of tumor that is highly associated with YAP activation, TM2 showed striking anti-proliferation efficacy as a single agent, which is consistent with the fact that therapeutic effects of TEAD inhibitors are mainly limited to mesothelioma models (Kaneda et al., 2020; Tracy T Tang et al., 2021). In colorectal cancer HCT116 and DLD I, single treatment of TM2 was insufficient to inhibit their growth, although they are also reported to be dependent on YAP activities. This might be interpreted by the activation of other oncogenic signaling pathways in these cancers, including Ras-MAPK activations. Indeed, YAP has been shown to converge with KRAS and can rescue cell viability induced by KRAS suppression (Shao et al., 2014), suggesting inhibiting YAP activities might be also rescued by other oncogenes. Consistently, significant synergy effects were observed when combining TM2 with a MEK inhibitor. These encouraging results suggested that rationalized combination of TEAD inhibitors with other inhibitors could significantly expand the utilities. In summary, our study disclosed TM2 as a promising new starting point for developing novel antitumor therapeutics against TEAD-YAP activities.

## Materials and methods

### Inhibition ofTEAD2 and TEAD4 auto-palmitoylation *In viiro*

Recombinant 6xHis-TEAD protein was treated with Compounds under indicated concentrations in 50 mM MES buffer (PH 6.4) for 30 mins. After incubation with 1 11M of alkyne palmitoyl-CoA (15968, Cayman) for 1 h, 50 11L of sample mixture was treated with 5 11L of freshly prepared “click” mixture containing 100 uM TBTA (678937, Sigma-Aldrich), 1 mM TCEP (C4706, Sigma-Aldrich), 1 mM CuS0_4_ (496130, Sigma-Aldrich), 100 uM Biotin-Azide (1167-5, Click Chemistry Tools) and incubated for another 1 h. The samples were then added 11 11L of 6xSDS loading buffer (BP-111R, Boston BioProducts) and denatured at 95°C for 5 mins. SDS-PAGE was used to analyze the samples. Palmitoylation signal was detected by streptavidin-HRP antibody (1:3000, S911, Invitrogen). The total protein level was detected by primary anti-His-tag antibody (1:10000, MA1-21315, Invitrogen) and secondary anti-mouse antibodies (1:5000, 7076S, Cell Signaling). The band intensities were quantified with ImageJ. The inhibition of auto-palmitoylation by compounds were normalized to DMSO. The IC50 curves were plotted with GraphPad prism6.

### Cell culture

Human H226, MST0-211H, H2052, H28, HCT116, DLD1 cells were obtained from ATCC (Manassas, VA). HEK293A, HCT116, DLD1 cells were cultured in Dulbecco’s modified Eagles media (DMEM) (Life Technologies) supplemented with 10% (v/v) fetal bovine serum (FBS) (Thermo/Hyclone, Waltham, MA), 100 units/mL penicillin and 100 11g/mL streptomycin (Life technologies) at 37°C with 5% C02. H226, MST0-211H, H2052, H28 cells were cultured in RPMI 1640 medium (Life technologies) supplemented with 10% (v/v) fetal bovine serum (FBS) (Thermo/Hyclone, Waltham, MA), 100 units/mL penicillin, 100 11g/mL streptomycin (Life technologies), 2.5g/L glucose and lmM sodium pyruvate at 37°C with 5% C02.

### Transfection

HEK293A cells was seed in 6 cm dishes overnight and transfected with plasmids using PEl reagent (lflg/flL). Briefly, PRK5-Myc-TEAD1 (33109, Addgene) and PEI were diluted in serum-free DMEM medium in two tubes (DNA: PEl ratio l:2). After standing still for 5 mins, mix them well and stay for another 20 mins. The mixture was then added to dishes directly.

### Inhibition ofTEAD palmitoylation in HEK293A cells

HEK293A cells with or without TEAD overexpression was pretreated with DMSO or TM2 in medium with 10% dialyzed fetal bovine serum (DFBS) for 8 hand labeled by Alkynyl Palmitic acid (1165, Click Chemistry Tools) for another 16 h. The cells were then washed and harvested by cold DPBS (14190250, Life Technologies). The cell pellets were isolated by centrifugation (500 x g, 10 min) and lysed by TEA lysis buffer (50mM TEA-HCl, pH 7.4, 150 mM NaCl, 1% Triton X-100, 0.2% SDS, lXProtease inhibitor-EDTA free cocktail (05892791001, Roche), phosphatase inhibitor cocktail (P0044, Sigma-Aldrich)) on ice for 30 mins. The protein concentration is determined using Bio-Rad assay and adjusted to 1 mg/mL. 100 11L of protein sample mixture was treated with 10 11L of freshly prepared “click” mixture containing 1 mM TBTA, 10 mM TCEP, 10 mM CuS0_4,_ 1 mM TBTA Biotin-Azide and incubated for 1 h at room temperature. The proteins were precipitated by chloroform/methanol/H_2_0 mixture and redissolved with 2%SDS in 0.1% PEST. The solution was diluted with 0.1% PEST and incubated with prewashed streptavidin agarose beads (69203-3, E M D MILLIPORE). After rotation at room temperature for 2 h, the beads were then pelleted by centrifugation (500 x g, 3 min) and washed with 0.2% SDS in PBS (3 x 1 mL). The bound proteins were eluted with a buffer containing 10 mM EDTA pH 8.2 and 95% formamide and analyzed with SDS-PAGE. Anti-Myc (1:1000, 2278S, Cell Signaling) or anti-pan-TEAD (1:1000, 13295, Cell Signaling) antibody were used to detect Myc-TEADl or pan-TEAD, respectively. Secondary antibody was anti-rabbit (1:5000, 7074S, Cell Signaling).

### Protein purification, crystallization, and structure determination

The recombinant human TEAD2 (residues 217-447, TEAD2 217-447) protein was purified and crystallized as described previously (Li et al., 2020b). Single crystals were soaked overnight at 20 °C with 5 mM TM2, 5% DMSO in reservoir solution supplemented with 25% glycerol and flashed-cooled in liquid nitrogen. Diffraction data was collected at beamline 19-ID (SBC-XSD) at the Advanced Photon Source (Argonne National Laboratory) and processed with HKL3000 program (Otwinowski and Minor, 1997). Best crystals diffracted 2.40 Å and exhibited the symmetry of space group C2 with cell dimensions of a124.1 A, b 62.3 A, c 79.9 A and 117.7°. Using TEAD2 structure (PDB ID: 3Ll5) as searching model, initial density map and model were generated by molecular replacement with Phaser in PHENIX (Adams et al., 2010). There are two TEAD2 molecules in the asymmetric unit. One TM2 molecule was built in the cavity of each TEAD2 molecule, and the remaining residues were manually built in COOT39 and refined in PHENIX. The final model (Rwork 0.184, Rfree 0.235) contains 400 residues, 30 water molecules and two TM2 molecules. Statistics for data collection and structure refinement are summarized in Table 1. The structure has been validated by wwPDB.40 Atomic coordinates and structure factors have been deposited to the Protein Data Bank under code 8CUH. Structural analysis and generation of graphics were carried out in PyMOL.

### Co-immunoprecipitation (Co-IP) assay

H226 cells were treated with DMSO or TM2 for 24 h. The cells were then washed and harvested by cold DPBS. The cell pellets were isolated by centrifugation (500 x g, 10 min) and lysed by lysis buffer (50mM Tris-HCl pH 7.5, 10% Glycerol, 1% NP-40, 300mM NaCl, 150mM KCl, 5mM EDTA, phosphatase inhibitor cocktail, complete EDTA-free protease inhibitors cocktail) on ice. After diluted with 50mM Tris-HCl pH 7.5, 10% Glycerol, 1% NP-40, 5mM EDTA, the protein samples were incubated with mouse anti-YAP antibody (sc-101199, Santa Cruz) overnight at 4°C and immunoprecipitated with prewashed protein A/G beads (P5030-1, UBPBio) for another 4 hat 4°C. The bound proteins were washed with 0.1% PEST for three times and eluted with 1xSDS loading buffer and analyzed with SDS-PAGE. Anti-TEAD1 (1:1000, 12292S, Cell Signaling), anti-pan-TEAD (1:1000, 13295, Cell Signaling) or anti-YAP (1:1000, 140745, Cell Signaling) antibody were used to detect TEAD1, pan-TEAD or YAP, respectively. Secondary antibody was anti-rabbit (1:5000, 7074S, Cell Signaling).

### Quantitative RT-PCR

H226 cells were treated with DMSO or TM2 for 24 h and used to extract RNA using the RNeasy mini kit (74104, Qiagen). The high-capacity cDNA reverse transcription kit (4368814, Life Technologies) was employed to obtain cDNA. Target genes expression *(Cyr61, CTGF andANKRDJ)* was measured with PowerUp SYB Green Master Mix kit (A25777, Life Technologies). *fJ-actin* was used as reference gene. The primers are shown below:

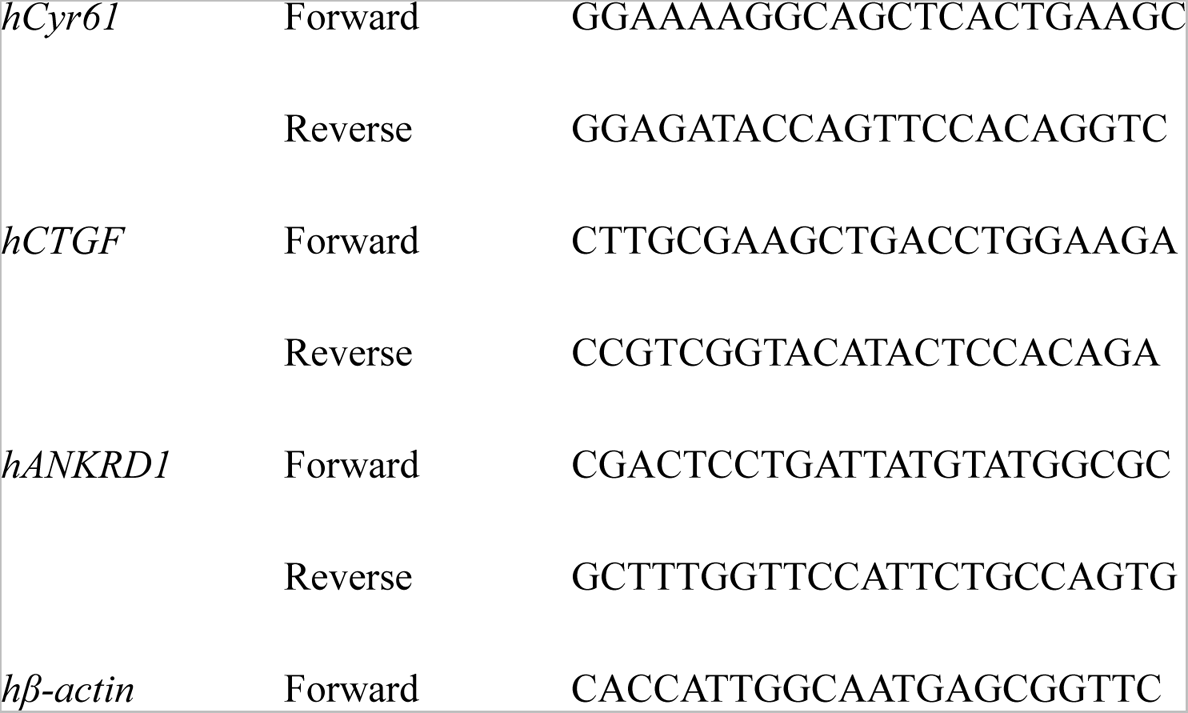
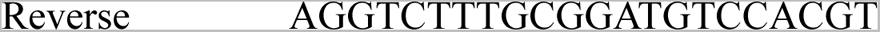

### RNA-seq analysis

The NCI-H226 cells were treated with TM2 at 1 11M for 24 hours. Total RNA was isolated with RNeasy Mini Kit (74104, Qiagen). The integrity of isolated RNA was analyzed using Bioanalyzer (Agilent Technologies). and the RNA-seq libraries were made by Novogene. All libraries have at least 50 million reads sequenced (150bp paired-end). The heatmap were generated using different expressed genes from TM2 treatment in NCI-H226 cells with Motpheus (https://software.broadinstitute.org/morpheus/). Principle component analysis (PCA) was determined by PCA function in M3C package in R. Gene Set Enrichment Analysis (GSEA) was performed using GSEA software from Broad Institute (http://software.broadinstitute.org/gsea/ index.jsp). The YAP_TAZ-TEAD Direct Target Genes set were generated with the published YAP/TAZ-TEAD target genes (Zanconato et al., 2015).

### Cell proliferation assay

H226, MST0-211H, H2052, H28, HCT116 and DLD1 cells were seed at a concentration of 500-2000 cells/well in 100 uL of culture medium in 96 well plates overnight and treated compounds with 3-fold dilutions of concentrations from 10 11M for 57 days. After removal of medium, each well was added 60 11L of MTT reagent (3-(4,5-dimethylthiazol-2-yl)-2,5-diphenyltetrazolium bromide) followed by incubation under 37°C for 4 h. The absorbance was measured by PerkinElmer EnVision plate reader.

### Drug Combination

The drug combination experiments were preformed using a drug combination matrix across 5 doses of TM2 (5 11M, 3-fold dilution) and 9 doses of Trametinib (10 11M, 3-fold dilution) in different tumor cell lines. Cell viability was determined at day 5 after the drugs administration by MTT. Drug synergy score was calculated followed Bliss rule. Synergy Score and Plot was generated by “Synergyfinder” package in R language.

### Organoids viability

Mouse hepatic progenitor organoids (70932, STEMCELL Tech) were seeded in 96 well plate using 20ul Matrigel (Corning, #354230) and cultured in HepatiCult™ Organoid Growth Medium (06031, STEMCELL Tech) with or without TM2. Medium was replaced after every 48 h with fresh compound. Organoid viability was measured by PrestoBlue™ HS Cell Viability Reagent (ThermoFisher, # P50200) following the manufacturer’s protocol.

### Immunofluorescence staining

Organoids were plated in 8 well chamber slide and fixed in 4% paraformaldehyde at 4°C for 1h. After permeabilization in 0.5% PEST, organoids were blocked with 2% BSA for 2 h and incubated with primary antibody overnight at 40C. Imaging was performed on Nikon A1RHD25 confocal microscope.

### Statistics

Data was analyzed by GraphPad prism6 and shown as mean ± SEM. All the biochemical experiments are repeated for at least 3 times and shown by representative images. Two-tailed t-test was used for P value calculation.

### Synthesis of TEAD inhibitors

All commercially available reagents were used without further purification. All solvents such as ethyl acetate, DMSO and Dichloromethane (DCM), were ordered from Fisher Scientific and Sigma-Aldrich and used as received. Unless otherwise stated, all reactions are conducted under air. Analytical thin­ layer chromatography (TLC) plates from Sigma were used to monitor reactions. Flash column chromatography was employed for purification and performed on silica gel (230-400 mesh). ^1^H NMR were recorded at 500 MHZ on JEOL spectrometer. ^13^C NMR were recorded at 125 MHZ on JEOL spectrometer. The chemical shifts were determined with residual solvent as internal standard and reported in parts per million (ppm).

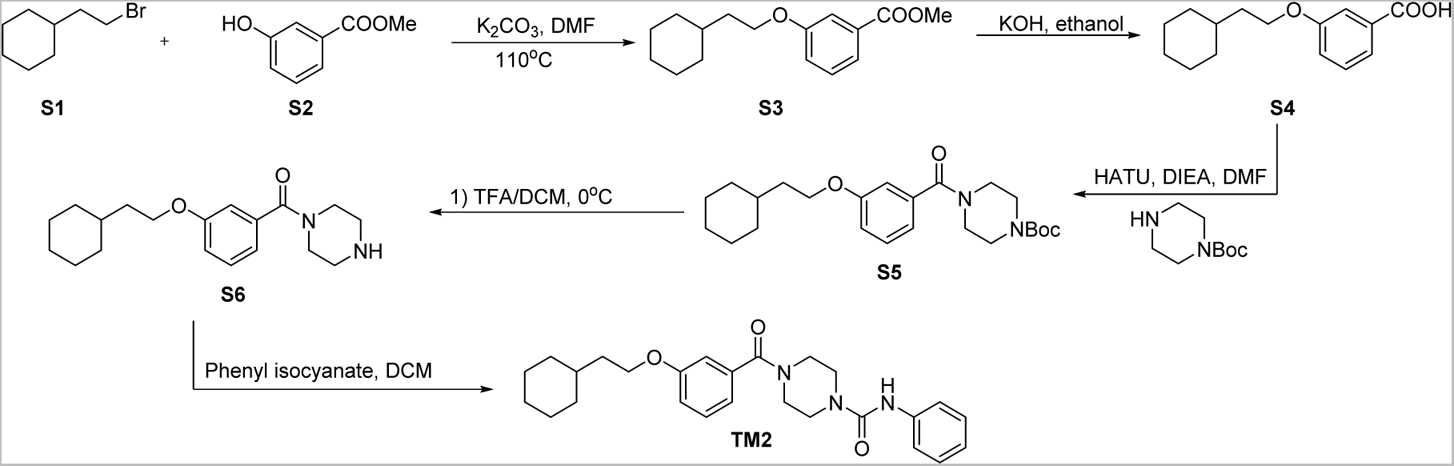

### Methyl3-(2-cyclohexylethoxy)benzoate (S3)

To a solution of methyl 3-hydroxybenzoate S2 (500 mg, 3.29 mmol) in DMF (7 mL) was added (2-bromoethyl)cyclohexane S1 (628.8 mg, 3.29 mmol) and K _2_C0_3_ (628.1 mg, 4.94 mmol). The mixture was then stirred at ll 0°C for 4 h. After cooling to temperature, the reaction mixture was diluted with water and extracted with Ethyl acetate. The combined organic layer was washed with brine, dried over Na_2_ S0_4_ and concentrated *in vacuo*. The crude residue was purified through silica gel chromatography to give S3 as colorless oil (780 mg, 90%). ^1^H NMR (500 MHz, Chloroform-d) *i5* 7.61 (d, *J* = 7.6 Hz, lH), 7.55 (t, *J* = 2.1 Hz, 1H), 7.33 (t, *J* = 7.9 Hz, 1H), 7.09 (dd, *J* = 8.2, 2.6 Hz, 1H), 4.03 (t, *J* = 6. 7Hz, 2H), 3.91 (s, 3H), 1.83-1.63 (m, 7H), 1.51 (ttt, *J* = 10.5, 6.8, 3.5 Hz, 1H), 1.33-l.ll (m, 3H), 0.98 (qd, *J* 11.9, 3.3 Hz, 2H).

### 3-(2-Cyclohexylethoxy)benzoic acid (S4)

To a solution of S3 (780 mg, 2.97 mmol) in ethanol (10 mL) was added saturated aqueous KOH (417 11L). The mixture was then stirred at room temperature overnight. After completion, the reaction was quenched with 1 N HCl on ice until PH was adjusted to 1. The mixture was then diluted with water and extracted with ethyl acetate. The combined organic layer was washed with brine, dried over anhydrous Na_2_ S0_4_ and concentrated *in vacuo* to give S4 (650 mg, 88%) which were used directly without further purification.

### *tert-*Butyl 4-(3-(2-cyclohexylethoxy)benzoyl)piperazine-1-carboxylate (S5)

To a solution of S4 (600 mg, 2.42 mmol) in DMF (20 mL) was added HATU (1.38 g, 3.63 mmol) and DIEA (862 flL, 4.84 mmol). After stirred for 5 mins, the solution was then added *tert-butyl* piperazine-1-carboxylate (450.6 mg, 2.42 mmol) and continuously stirred at room temperature overnight. After completion, the reaction was quenched with water and extracted with ethyl acetate. The combined organic layer was washed with 1 N HCl, saturated NaHC0_3_, brine, dried over anhydrous Na_2_ S0_4_ and concentrated *in vacuo*. The crude residue was purified through silica gel chromatography to give S5 as a white solid (950 mg, 94%). ^1^H NMR (500 MHz, Chloroform-d) *δ* 7.30 (t, *J* = 8.0 Hz, 1H), 6.96-6.89 (m, 3H), 3.99 (t, *J* = 6.7 Hz, 2H), 3.82-3.31 (m, 8H), 1.79-1.62 (m, 7H), 1.54-1.39 (m, 1H) 1.47 (s, 9H), 1.32-1.10 (m, 3H), 0.96 *(qd,J*11.9, 3.0 Hz, 2H).

### (3-(2-Cyclohexy1ethoxy)pheny1)(piperazin-1-y1)methanone (S6)

To a solution of S5 (890 mg, 2.13 mmol) in DCM (4 mL) was added trifluoroacetic acid (4 mL) dropwise on ice. The mixture was continuously stirred on ice for 30 mins. After completion, the reaction was quenched with saturated NaHC0_3_ dropwise on ice. The mixture was then diluted with water and extracted with ethyl acetate. The combined organic layer was washed with brine, dried over anhydrous Na_2_ S0_4_ and concentrated *in vacuo* to give S6 which were used directly without further purification.

### 4-(3-(2-cyclohexy1ethoxy)benzoy1)-N-pheny1piperazine-1-carboxamide (TM2)

To a solution of S6 (100 mg, 0.403 mmol) in DCM (4 mL) was added isocyanate phenyl isocyanate (63.1 flL, 0.484 mmol). The reaction mixture was stirred at room temperature for 2 h. The reaction was quenched with water and extracted with DCM. The combined organic layer was washed with brine, dried over anhydrous Na_2_ S0_4_ and concentrated in vacuo. The crude residue was purified through silica gel chromatography to give TM2 as a white solid (160 mg, 91%). ^1^H NMR (500 MHz, Chloroform-d) o 7.36-7.24 (m, 5H), 7.04 (t, *J* = 7.3 Hz, 1H), 6.98-6.89 (m, 3H), 6.77 (brs, 1H), 3.99 (t, *J* = 6.7 Hz, 2H), 3.93-3.35 (m, 8H), 1.78-1.62 (m, 7H), 1.54-1.44 (m, 1H), 1.30-1.12 (m, 3H), 0.97 (qd, J = 12.1, 2.9 Hz, 2H). 13C NMR (125 MHz, Chloroform-d) o 170.62, 159.45, 155.21, 138.85, 136.47, 129.85, 129.02, 123.55, 120.41, 118.87, 116.46, 113.17, 66.28, 47.46 (brs), 44.22, 42.01 (brs), 36.64, 34.61, 33.39, 26.60, 26.33.

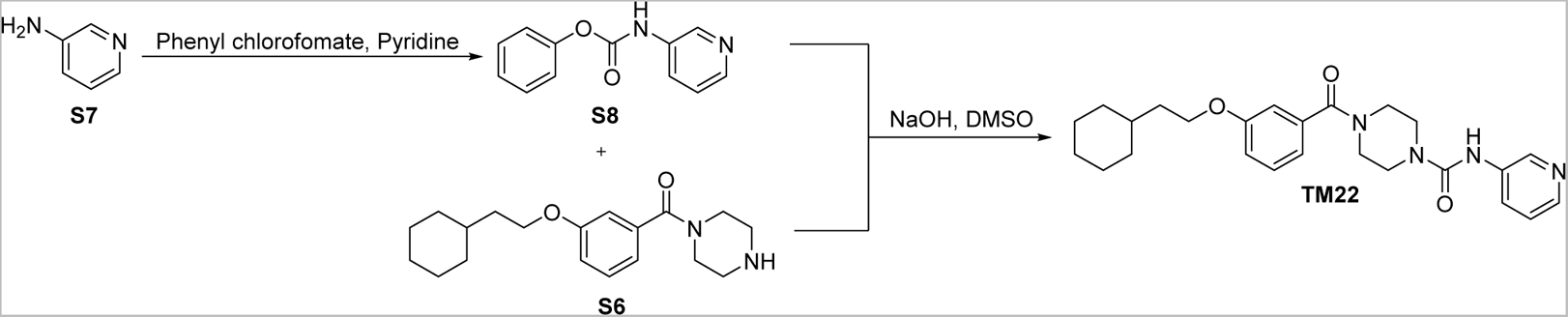

### Phenyl pyridin-3-ylcarbamate (S8)

To a solution of Pyridin-3-amine S7 (188.2 mg, 2 mmol) in pyridine (5 mL) was added phenyl chloroformate (274 flL, 2.2 mmol). The reaction mixture was stirred at room temperature overnight. The mixture was quenched by the addition of ethyl acetate and 10% critic acid. The organic layer was washed with saturated NaHC0_3_, brine, dried over Na_2_ S0_4._ The organic solvents were removed in vacuo to give carbamate S8 which was used directly for the next step.

### 4-(3-(2-cyclohexylethoxy)benzoyl)-N-(pyridin-3-yl)piperazine-1-carboxamide (TM22)

To a solution of S6 (30 mg, 0.095 mmol) in DMSO (1 mL) was added carbamate (40.7 mg, 0.19 mmol) and NaOH (114 flL, 0.114 mmol, 10 N). The reaction mixture was stirred at room temperature for 2 h. The reaction was quenched with water and extracted with ethyl acetate. The combined organic layer was washed with brine, dried over anhydrous Na_2_ S0_4_ and concentrated *in vacuo*. The crude residue was purified through silica gel chromatography to give TM22 as a white solid (36.1 mg, 87%). ^1^H NMR (500 MHz, Chloroform-d) *δ* 8.46 (d, *J* = 2.6 Hz, 1H), 8.26 (dd, *J* = 4.8, 1.4 Hz, 1H), 7.96 (dt, *J* 8.4, 2.1 Hz, 1H), 7.34-7.20 (m, 3H), 6.99-6.87 (m, 3H), 3.99 (t, *J* = 6.7 Hz, 2H), 3.88-3.37 (m, 8H), 1.77-1.63 (m, 7H), 1.55-1.45 (m, 1H), 1.29-1.13 (m, 3H), 0.96 (qd, *J* = 12.0, 2.9 Hz, 2H). ^13^C NMR (125 MHz, Chloroform-d) *δ* 170.70, 159.50, 155.05, 144.25, 141.49, 136.36, 136.25, 129.93, 127.78, 123.78, 118.82, 116.49, 113.21, 66.32, 47.43 (brs), 44.24, 42.00 (brs), 36.65, 34.64, 33.41, 26.62, 26.35.

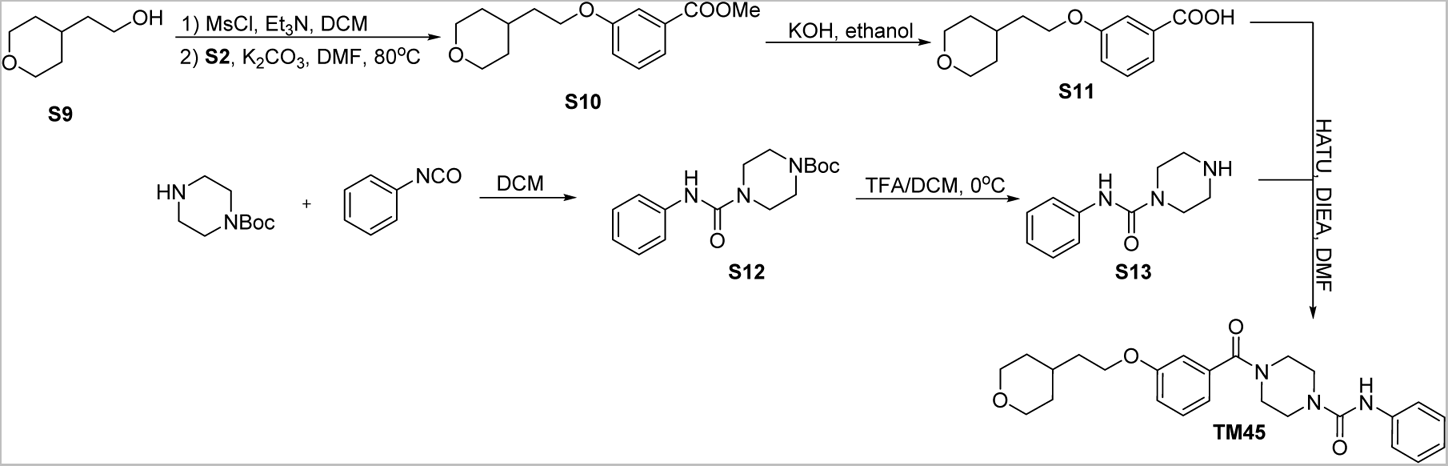

### Methyl3-(2-(tetrahydro-2H-pyran-4-yl)ethoxy)benzoate (S10)

To a solution of S9 (400 mg, 3.07 mmol) in anhydrous DCM (20 mL) was added Et_3_N (642 flL, 4.61 mmol), MsCl (285 flL, 3.68 mmol) at 0°C. The solution was stirred at room temperature. After completion, the reaction mixture was diluted with water, extracted with DCM, washed with saturated aqueous NaHC0_3._ The combined organic layer was dried over anhydrous Na_2_ S0_4_ and concentrated *in vacuo* to give the methanesulfonate. The methanesulfonate was then dissolved in DMF (10 mL) followed by cautiously adding S2 (513.8 mg, 3.38 mmol) and K_2_C0_3_ (848.6 mg, 6.14 mmol). The resulting suspension was further stirred at 80°C for 4 h. The reaction mixture was extracted with ethyl acetate, then washed with water, brine. The organic phase was dried over anhydrous Na_2_ S0_4_ and concentrated *in vacuo*. The crude residue was purified through silica gel chromatography to give S10 as colorless oil (680 mg, 84%). ^1^H NMR (500 MHz, Chloroform-d) *i5* 7.62 (dd, *J* = 7.5, 1.3 Hz, 1H), 7.54 (t, *J* 2.1 Hz, 1H), 7.33 (t, *J* = 7.9 Hz, 1H), 7.08 (dd, *J* = 8.4, 2.6 Hz, 1H), 4.05 (t, *J* = 6.2 Hz, 2H), 3.97 (ddd, *J* = 11.4, 4.5, 1.7 Hz, 2H), 3.91 (s, 3H), 3.41 (td, *J* ll.8, 2.1 Hz, 2H), 1.85-1.73 (m, 3H), 1.67 *(dq,J*13.3, 2.0 Hz, 2H), *1.37(qd,J*11.9, 4.4Hz, 2H).

### 3-(2-(Tetrahydro-2H-pyran-4-yl)ethoxy)benzoic acid (S11)

S11 was prepared as described for **S4** (670 mg, 2.53 mmol) from **SlO** and were used directly without further purification.

### *tert-*Butyl 4-(phenylcarbamoyl)piperazine-1-carboxylate (S12)

Sl2 was prepared as described for TM2 from tert-butyl piperazine-1-carboxylate (1 g, 5.37 mmol) and phenyl isocyanate (767.5 mg, 6.44 mmol) as a white solid (quantitative). ^1^H NMR (500 MHz, Chloroform-d) *δ* 7.35 (d, *J* = 7.6 Hz, 2H), 7.29 (td, *J* = 8.5, 8.0, 2.3 Hz, 2H), 7.08-7.02 (m, 1H), 6.37 (s, 1H), 3.49 (s, 8H), 1.49 (s, 9H).

### N-phenylpiperazine-1-carboxamide (S13)

S13 was prepared as described for S6 (800 mg, 2.62 mmol) from S12 and were used directly without further purification.

### *N-*phenyl-4-(3-(2-(tetrahydro-2H-pyran-4-yl)ethoxy)benzoyl)piperazine-1-carboxamide (TM45)

TM45 was prepared as described for S5 from S11 (40 mg, 0.16 mmol) and S13 (39.4 mg, 0.192 mmol) as a white solid (44 mg, 63%). ^1^H NMR (500 MHz, Chloroform-d) *δ* 7.35-7.29 (m, 3H), 7.29-7.24 (m, 2H), 7.07-7.01 (m, 1H), 6.98-6.89 (m, 3H), 6.73 (s, 1H), 4.01 (t, *J* = 6.1 Hz, 2H), 3.96 (dd, *J* = 11.1, 3.6, 2H), 3.86-3.43 (m, 8H), 3.39 (td, *J* = 11.8, 2.0 Hz, 2H), 1.81-1.71 (m, 3H), 1.68-1.61 (m, 2H), 1.40-1.30 (m, 2H). ^13^C NMR (125 MHz, Chloroform-d) *δ* 170.52, 159.30, 155.19, 138.85, 136.56, 129.88, 129.01, 123.55, 120.37, 119.03, 116.40, 113.17, 68.06, 65.48, 47.43 (brs), 44.21, 42.01 (brs), 36.15, 33.07, 32.01.

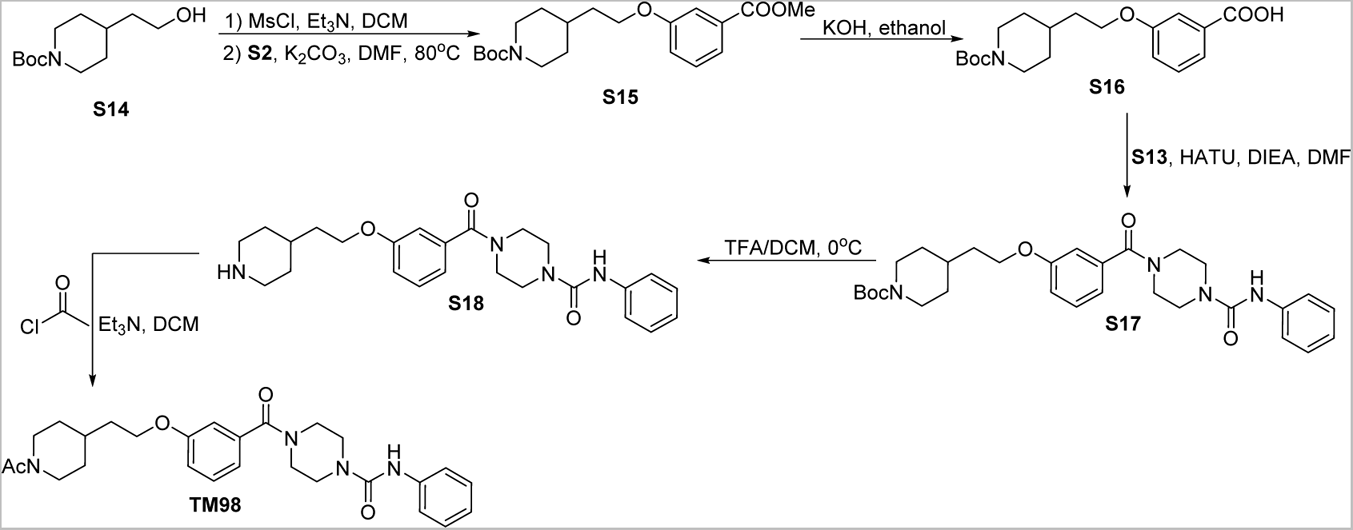

### *tert-*Butyl 4-(2-(3-(methoxycarbonyl)phenoxy)ethyl)piperidine-1-carboxylate (S15)

Sl5 was prepared as described for SlO from Sl4 (480 mg, 2.09 mmol) and S12 (318 mg, 2.09 mmol) as a white solid (530 mg, 70%). ^1^H NMR (500 MHz, Chloroform-d) *δ* 7.62 (dd, *J* = 7.7, 1.3 Hz, 1H), 7.54 (dd, *J* = 2.7, 1.3 Hz, 1H), 7.33 (t, *J* = 7.9 Hz, 1H), 7.08 (ddd, *J* = 8.3, 2.6, 1.2 Hz, 1H), 4.19-4.05 (m, 2H), 4.05 (t, *J* = 6.1 Hz, 2H), 3.91 (s, 3H), 2.80-2.64 (s, 2H), 1.80-1.65 (m, 5H), 1.46 (s, 9H), 1.23-1.11 (m, 2H).

### 3-(2-(1-(tert-Butoxycarbonyl)piperidin-4-yl)ethoxy)benzoic acid (S16)

S16 was prepared as described for S4 from S15 (380 mg, 1.05 mmol) and were used directly without further purification.

### tert-Butyl 4-(2-(3-(4-(phenylcarbamoyl)piperazine-1-carbonyl)phenoxy)ethyl)piperidine-1-carboxylate (S17)

S17 was prepared as described for S5 from S16 (200 mg, 0.572 mmol) and S13 (140.9 mg, 0.686 mmol) as a white solid (270 mg, 88%). ^1^H NMR (500 MHz, Chloroform-d) *δ* 7.33 (t, *J* = 7.9 Hz, 3H), 7.30-7.24 (m, 2H), 7.03 (tt, *J* = 7.4, 1.3 Hz, 1H), 6.98-6.89 (m, 3H), 6. 78 (brs, 1H), 4.16-4.04 (m, 2H), 4.00 (t, *J* 6.2 Hz, 2H), 3.87-3.35 (m, 8H), 2.70 (s, 2H), 1.82-1.64 (m, 5H), 1.45 (s, 9H), 1.21-1.11 (m, 2H).

### *N-*phenyl-4-(3-(2-(piperidin-4-yl)ethoxy)benzoyl)piperazine-1-carboxamide (S18)

S18 was prepared as described for S6 from S17 (175mg, 0.33 mmol) and were used directly without further purification.

### 4-(3-(2-(1-acetylpiperidin-4-yl)ethoxy)benzoyl)-N-phenylpiperazine-1-carboxamide (TM98)

S18 (25 mg, 0.0573 mmol) was then dissolved in DCM (1.5 mL). The solution was added Et3N (16 flL, 0.115 mmol) and acetyl chloride (4.9 flL, 0.0688 mmol) on ice. The reaction mixture was stirred at room temperature for 2 h. After completion, the reaction was quenched with saturated NaHC0_3_ and extracted with ethyl acetate. The combined organic layer was washed with brine, dried over anhydrous Na_2_ S0_4_ S18 as a colorless oil (20 mg, 73%). ^1^H NMR (500 MHz, Chloroform-d) *i5* 7.38-7.24 (m, 5H), 7.04 (t, *J* 7.3 Hz, 1H), 6.94 (ddt, *J* = 10.3, 6.1, 2.5 Hz, 3H), 6.86-6.75 (m, 1H), 4.60 (d, *J* = 13.1 Hz, 1H), 4.02 (t, *J* 5.9 Hz, 2H), 3.92-3.36 (m, 9H), 3.04 (t, *J* 13.0 Hz, 1H), 2.54 (t, *J* 13.0 Hz, 1H), 2.08 (s, 3H), 1.84-1.71 (m, 5H), 1.27-1.10 (m, 2H). ^13^C NMR (125 MHz, Chloroform-d) *i5* 170.50, 168.98, 159.17, 155.24, 138.92, 136.64, 129.92, 129.02, 123.52, 120.34, 119.17, 116.37, 113.28, 65.57, 46.77, 44.26, 41.90, 35.57, 33.20, 32.78, 31.83, 21.61. HRMS (ESI): calcd for C27H35N404 [M+Ht, 479.2658; found, 479.2653.

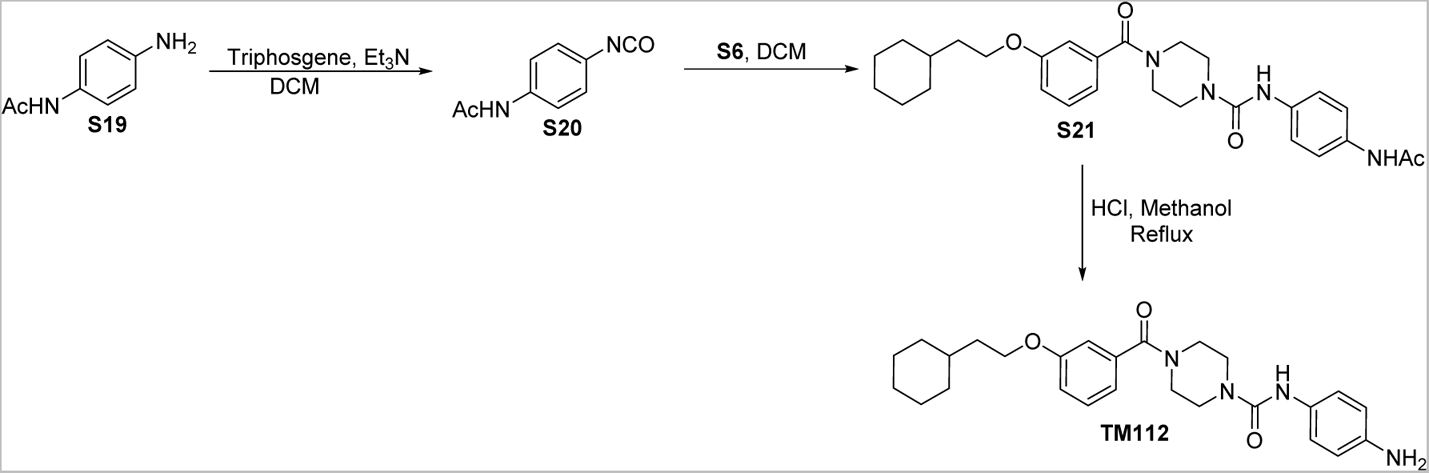

### *N-(* 4-isocyanatophenyl)acetamide (S20)

To a solution oftriphosgene (311.6 mg, 1.05 mmol) in DCM (6 mL) was added a solution of Et_3_N (0.9 mL, 6.45 mmol) and S19 (450.5 mg, 3 mmol) in DCM (6 mL) dropwise on ice. The mixture was continuously stirred at rt for 1h. The reaction was quenched with saturated NaHC0_3_ dropwise on ice. The mixture was then diluted with water and extracted with ethyl acetate. The combined organic layer was washed with brine, dried over anhydrous Na_2_ S0_4_ and concentrated in vacuo to give S20 which were used directly without further purification.

### *N-(* 4-Acetamidophenyl)-4-(3-(2-cyclohexylethoxy)benzoyl)piperazine-1-carboxamide (S21)

S21 was prepared as described for TM2 from S6 (120 mg, 0.376 mmol) and *N-(3-*isocyanatophenyl)acetamide (79.5 mg, 0.451 mmol) as a white solid (100.5 mg, 54%). ^1^H NMR (500 MHz, Chloroform-d) *i5* 7.40 (d, *J* = 8.4 Hz, 2H), 7.33-7.26 (m, 3H), 7.15 (brs, 1H), 6.98-6.89 (m, 3H), 6.39 (brs, 1H), 3.99 (t, *J* = 6.7 Hz, 3H), 3.92-3.35 (m, 8H), 2.15 (s, 3H), 1.77-1.62 (m, 7H), 1.53-1.44 (m, 1H), 1.29-1.10 (m, 3H), 1.01-0.90 (m, 2H).

### *N-(* 4-aminophenyl)-4-(3-(2-cyclohexylethoxy)benzoyl)piperazine-1-carboxamide (TM112)

To a solution of S21 (80 mg, 0.161 mmol) in methanol (2 mL) was added 2 N HCl (4 mL). The reaction was refluxed for 2 h. After cooling down to rt, the reaction mixture was basified with saturated NaHC0_3_ on ice and extracted with ethyl acetate. The combined organic layer was washed with brine, dried over anhydrous Na_2_ S0_4_ and concentrated *in vacuo*. The crude residue was purified through silica gel chromatography to give TM112 as colorless oil (23.8 mg, 33%). ^1^H NMR (500 MHz, Chloroform-d) i5 7.31 (t, *J* = 8.0 Hz, 1H), 7.09 (d, *J* = 8.6 Hz, 2H), 6.98-6.90 (m, 3H), 6.63 (d, *J* = 8.6 Hz, 2H), 6.23 (s, lH), 4.00 (t, *J* = 6.7 Hz, 2H), 3.93-3.26 (m, lOH), 1.78-1.64 (m, 7H), 1.55-1.45 (m, 1H), 1.31-1.15 (m, 3H), 0.97 (qd, *J* = 12.1, 3.0 Hz, 2H). ^13^C NMR (125 MHz, Chloroform-d) *i5* 170.63, 159.47, 155.85, 143.21, 136.61, 129.85, 129.80, 123.29, 118.97, 116.49, 115.70, 113.21, 66.32, 44.26, 36.68, 34.66, 33.42, 26.64, 26.37.

## Additional information

### Author contributions

Lu Hu, Conceptualization, Data Curation, Formal analysis, Investigation, Writing-original draft, Writing-review and editing; Sun Yang, Data Curation, Formal analysis, Investigation, Writing-review and editing; Shun Liu, Data Curation, Formal analysis, Investigation; Hannah Erb, Data Curation, Formal analysis, Investigation; Xuelian Luo, Supervision, Formal analysis, Writing-review and editing; Xu Wu, Conceptualization, Supervision, Resources, Funding acquisition, Formal analysis, Writing-original draft, Writing-review and editing.

## Additional files

### Data availability

The crystal structure of TEAD2 YBD in complex with TM2 has been deposited in the Protein Data Bank with accessiOn codes 8CUH. The raw RNA-seq data of H226 treated with TM2 has been deposited _1U_ NCBI GEO DataSets and _IS_ accessible at https://www.ncbi.nlm.nih.govIgeo/querya!cc.cgi?acc=GSE203421.

The following dataset was generated:

| Authors | Year | Dataset title | Dataset URL | Database and Identifier |
| --- | --- | --- | --- | --- |
| Xu Wu, Yang Sun, Lu Hu | 2022 | Next Generation Sequencing | https://www.ncbi.nlm.nih.gov/geo/query/acc.cgi?ac | NCBI GEO DataSets, GSE203421 |
|  |  | Quantitative Analysis<br>of TEAD<br>Palmitoylation<br>Inhibitor TM2 in NCI-H226 cells | c=GSE203421 |  |

The following previously published dataset was used:

| Author(s) | Year | Dataset title | Dataset URL | Database and<br>Identifier |
| --- | --- | --- | --- | --- |
| Calvet L, Dos-Santos O,<br>Spanakis E, Valence S,<br>Jean-Baptiste V, Le Bail<br>J, Buzy A, Paul P, Henry<br>C, Pollard J, Sidhu S,<br>Moll J, Debussche L,<br>Valtingojer I | 2022 | Human malignant<br>mesothelioma NCI-H226 cells treated<br>with TEAD inhibitor<br>K-975 in SCID mice | https://www.ncbi.nlm.nih.gov/geo/query/acc.cgi?acc=GSE196726 | NCBI GEO DataSets,<br>GSE196726 |

